# Joint distribution of nuclear and cytoplasmic mRNA levels in stochastic models of gene expression: analytical results and parameter inference

**DOI:** 10.1101/2024.04.29.591679

**Authors:** Yiling Wang, Juraj Szavits-Nossan, Zhixing Cao, Ramon Grima

## Abstract

Common stochastic models of gene expression predict the analytical distribution of the total mRNA level per cell but not at subcellular resolution. Here, for a wide class of models of transcription initiation, we obtain an exact steady-state solution for the joint distribution of nuclear and cytoplasmic mRNA levels per cell. Correcting the solution for extrinsic noise and fitting to single human cell data, we precisely quantify the extent of bursty expression in thousands of genes and associate it with their biological functions.

---

Gene expression is an inherently noisy process [1–4] resulting in significant variation of mRNA and protein numbers between cells. Stochastic models have been studied to shed light on the biophysical processes leading to this variability [5–7] and to fit data and thereby estimate the parameter values specific to each gene [8–11].

The vast majority of models of stochastic gene expression are based on the telegraph model whereby a gene is in one of two states (active and inactive), mRNA is produced from the active state and subsequently decays [12, 13]. The mRNA in this model has often been interpreted as the total mRNA in a cell. However, nowadays mRNA levels can be measured simultaneously in both the nucleus and the cytoplasm [14–17]. Hence, more refined stochastic models of gene expression are needed which can fit this type of data and to also understand how the spatial partitioning of molecular systems may play a role in the reduction of noise [18].

Models involving more than one type of mRNA, i.e. describing all or a subset of nascent, mature nuclear and mature cytoplasmic mRNA levels, have been constructed and solved either numerically [15, 19] or analytically for various special cases of interest [20–27]. The analytical solutions provide expressions for either the moments or the marginal distributions of mRNA levels, except Ref. [20] which additionally provides the joint distribution of nuclear and cytoplasmic mRNA. The model in Ref. [20] is a special case of the telegraph model in which the gene is active for an infinitesimally short time during which a random number of mRNAs (a burst) are produced. This model is problematic because the assumption of instantaneous bursting is debatable for eukaryotic genes since many of these spend many minutes in the active state [28]. As well, the assumption of exponentially distributed off times is in contrast to experimental evidence for some mammalian genes [29].

In this paper, we consider a general stochastic model of gene expression that does not make the aforementioned assumptions and derive the exact joint distribution for mRNA levels measured at two different points of the mRNA lifecycle —these two types of mRNA can be interpreted as nascent and mature nuclear mRNA or unspliced and spliced mRNA or mature nuclear and cytoplasmic mRNA. We verify the accuracy of the solution using Monte Carlo simulations and finally use it to devise an accurate and computationally efficient method for the inference of gene-specific kinetic parameters from commonly available data.

## General theory

We consider a general stochastic model of nuclear and cytoplasmic mRNA fluctuations in which the production of nuclear mRNA constitutes a renewal process [30], i.e. we assume that the interarrival times of successive nuclear mRNA production events are independent and identically distributed random variables. This assumption is supported by experimental evidence [31]. We denote the probability density function of the interarrival times by *f* (*t*) and assume that the mean interarrival time 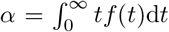 is finite. After a time delay *τ*, nuclear mRNAs (*N*) are then exported to the cytoplasm, thus becoming cytoplasmic mRNA (*M*). These molecules then degrade at a rate *d*. For now, we assume that *τ* is deterministic because this makes the analytic computations tractable. In accordance with previous work [32], we do not model the degradation of nuclear mRNA because bona fide coding transcripts are specifically protected from RNA decay enzymes in the nucleus [33]. The whole model can be summarised by the following reaction scheme,

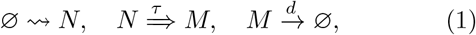

where ⇝ denotes a reaction that takes a random amount of time whose probability density function is *f* (*t*) and denotes a reaction of fixed duration *τ*. In Fig. 1 we illustrate a specific example of a gene expression model of the type of Eq. (1).

**FIG. 1.**
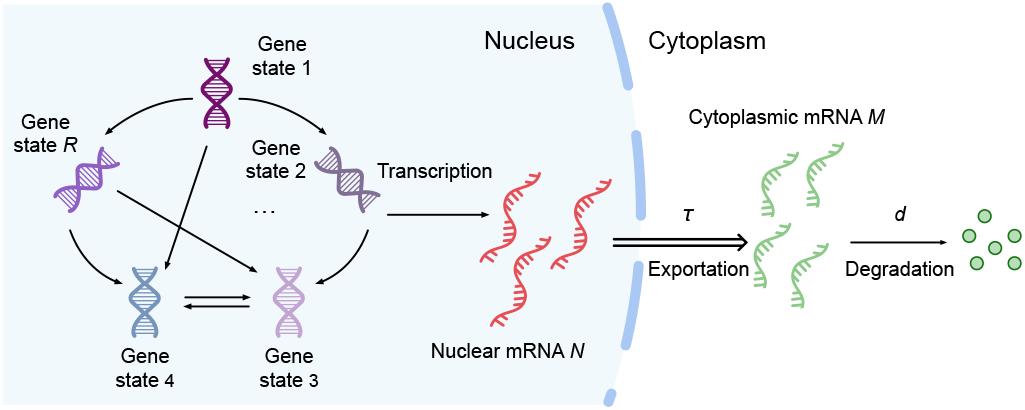
Illustration of a gene expression model describing initiation (gene switching between a finite number *R* of inactive and active states), transcription from an active state and export after a finite time *τ* of mRNA from the nucleus to the cytoplasm where it is finally degraded. This is an example of a wide class of systems in which production of nuclear mRNA is a renewal process; for such systems we derive the steady-state joint distribution of nuclear and cytoplasmic mRNA counts.

To compute the steady-state joint distribution of nuclear and cytoplasmic mRNA, we employ queueing theory, whose application to gene expression has been recently reviewed in Ref. [34]. Queueing theory is a mathematical theory that describes customers arriving to a facility where they receive a service and leave [35]. Queueing systems are typically described using Kendall’s short-hand notation *A/S/n*, where *A* denotes the arrival process, *S* the service process and *n* is the number of servers [36]. Eq. (1) is `mathematically` equivalent to a queueing system consisting of two queues connected in a series. The first queue describes the production (arrival) of nuclear mRNA (customers) and its processing (service) into cytoplasmic mRNA. It corresponds to the *G*/*D*/∞ queue, where *G* denotes renewal arrivals with general (arbitrary) interarrival time distribution and *D* denotes deterministic (fixed-time) service. The output of this queue feeds into the second queue, which describes the degradation (service) of cytoplasmic mRNA. The second queue corresponds to the *G*/*M*/∞ queue, where *M* denotes Markov or memoryless service. Since the nuclear processing is deterministic, both queues have the same interarrival time distribution. We assume that both types of mRNA are available for processing immediately after they are created, hence both queues have infinitely many servers.

Let *N* (*t*) and *M* (*t*) denote the number of nuclear and cytoplasmic mRNA at time *t*, respectively. We are interested in the joint distribution *P* (*n, m*) of *N* (*t*) = *n* and *M* (*t*) = *m* in the limit of *t*. → ∞ Let *ξ*(*t*) denote the residual time from *t* until the next cytoplasmic mRNA is produced, and *f*_*m*_(*x*) the probability density function of *ξ*(*t*) = *x* conditioned on *M* (*t*) = *m* in the stationary limit (i.e. when *t* → ∞). We note that because nuclear mRNA is processed deterministically, *N* (*t*) is equal to the number of nuclear mRNA arrivals between *t* − *τ* and *t*, of which the first arrival occurs at time *t* − *τ* + *ξ*(*t*). For *ξ*(*t*) ≤ *τ*, let *Y* (*τ* − *ξ*(*t*)) denote the number of nuclear RNA arrivals from *t* − *τ* + *ξ*(*t*) until *t*, not including the arrival at *t* − *τ* + *ξ*(*t*), and let *K*_*n*−1_(*τ − ξ*(*t*)) denote the probability of *Y* (*τ − ξ*(*t*)) = *n* − 1. From these definitions, it follows that the stationary joint distribution *P* (*n, m*) can be written as

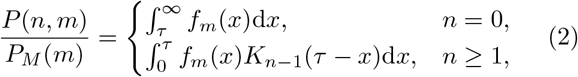

where *P*_*M*_ (*m*) is the stationary probability of *M* (*t*) = *m*. The interpretation of Eq. (2) is the following. The event of *N* (*t*) = *n* and *M* (*t*) = *m* is equivalent of having: (i) *ξ*(*t*) = *x* and *M* (*t*) = *m*, whose joint probability is *f*_*m*_(*x*)d*xP*_*M*_ (*m*), and (ii) *Y* (*τ* − *ξ*(*t*)) = *n* − 1, whose probability is *K*_*n*−1_(*τ* − *x*). The product of these two probabilities is then integrated over all *x* from 0 until *τ*. For *n* = 0, there are no arrivals between *t* − *τ* and *τ*, hence we integrate *f*_*m*_(*x*)d*xP*_*M*_ (*m*) over *x* ≥ *τ*.

In the context of queueing theory, *P*_*M*_ (*m*) is the stationary queue-length distribution of the *G*/*M*/∞ queue, which has been computed in Ref. [37] for an arbitrary interarrival time distribution with finite mean *α*. The probability *K*_*n*−1_(*τ*–*x*) is known from renewal theory [30] and reads

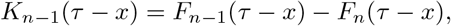

where 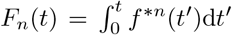 and *f* ^\**n*^ is the *n*-fold convolution of *f*. Finally, the probability density function *f*_*m*_(*x*) has been computed in Ref. [37] and reads

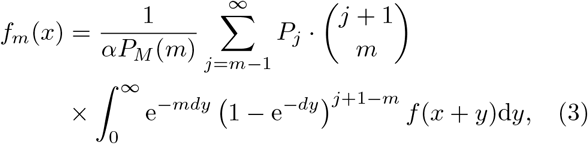

where *α* is the mean interarrival time and *P*_*j*_ is given by

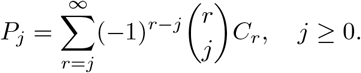

In the above expression, coefficients *C*_*r*_ are defined as

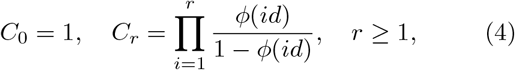

and 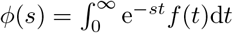.

The joint probability *P* (*n, m*) is most conveniently computed via the probability generating function (PGF) 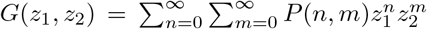 Using Eq. (2), *G*(*z*_1_, *z*_2_) can be compactly written as

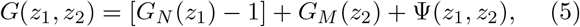

where *G*_*N*_ (*z*_1_) and *G*_*M*_ (*z*_2_) are the probability generating functions of nuclear and cytoplasmic mRNA distributions, respectively. The expression for *G*_*M*_ (*z*_2_) is known explicitly [37],

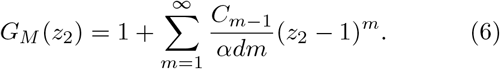

On the other hand, *G*_*N*_ (*z*_1_) [38] and Ψ(*z*_1_, *z*_2_) can be computed by inverting their Laplace transforms with respect to the nuclear processing time *τ*,

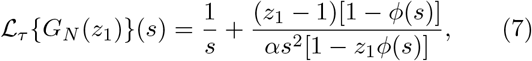

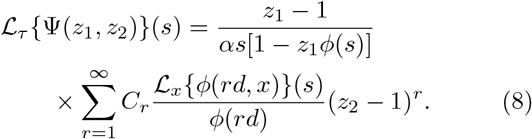

In the last expression, 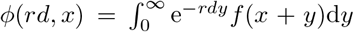 and ℒ _*x*_ *ϕ*(*rd, x*) (*s*) is the Laplace transform of *ϕ*(*rd, x*) with respect to *x*. The derivation of Eqs. (5) and (8) is presented in Supplemental Material (SM) I.

**Model I**. All results so far have been derived for an arbitrary (general) interarrival time distribution. To specify a particular model of nuclear mRNA production, we need to specify the form of *f* (*t*). Typically, nuclear mRNA production is modelled as a Markov process in which the gene switches between a number of discrete states and produces mRNA from one or more of these states [39]. The most widely used biophysical model is the telegraph model [12], whose two compartment extension is given by

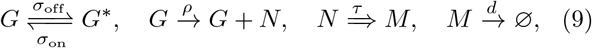

where *G*^*^ and *G* are the inactive and active gene states, respectively. Henceforth Eq. (9) will be referred to as

Model I. Applying the general result obtained using queueing theory (Eq. (5)) we obtain an analytical expression for *G*(*z*_1_, *z*_2_) of Model I (for details see SM II)

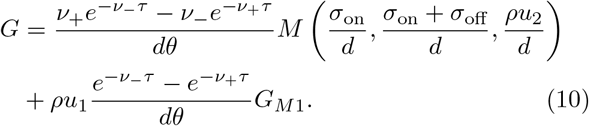

Here *G*_*M*1_ = *αM* (1 + *σ*_on_*/d*, 1 + *σ*_off_*/d* + *σ*_on_*/d, x*_2_) with *α* = *σ*_on_/(*σ*_off_ + *σ*_on_), *x*_2_ = *ρu*_2_*/d* and *M* (*a, b, x*) being the Kummer function. Additionally, 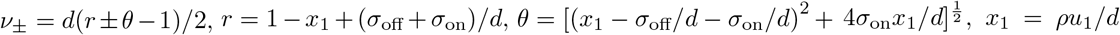 and *u*_1,2_ = *z*_1,2_ − 1. An alternative (non-queueing theory based) derivation of the solution from the delay master equation can be found in SM III. We note that the generating function marginalized to include only information about nuclear mRNA, i.e. *G*(*z*_1_, *z*_2_ = 1), has been previously derived [38, 40]; note that in these papers the species *N* is interpreted as the bound RNA polymerase rather than mature nuclear mRNA. Finally, the joint distribution solution is constructed using the relation

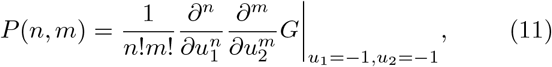

which can be efficiently calculated by a symbolic computation program. We emphasize that unlike current solutions of the joint distribution [20], our solution does not assume that the time spent in state *G* is infinitesimally short (instantaneous bursts).

In Fig. 2a upper panel, we use P-P plots (probability–probability plots) to show the accuracy of the exact solution given by Eqs. (10) and (11) by comparing it with the joint distributions 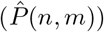 computed from 10^6^ independent runs of the delay stochastic simulation algorithm (delay SSA, the rejection method in Ref. [41]). In this plot, we show all the pairs of conditional cumulative probabilities 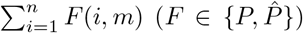 for any possible values of *n, m* ∈, [0, *N*_max_], where *N*_max_ is the largest mRNA count recorded in the SSA simulations. All the points fall on the line *y* = *x*, thereby verifying the accuracy of the analytical solution.

**FIG. 2.**
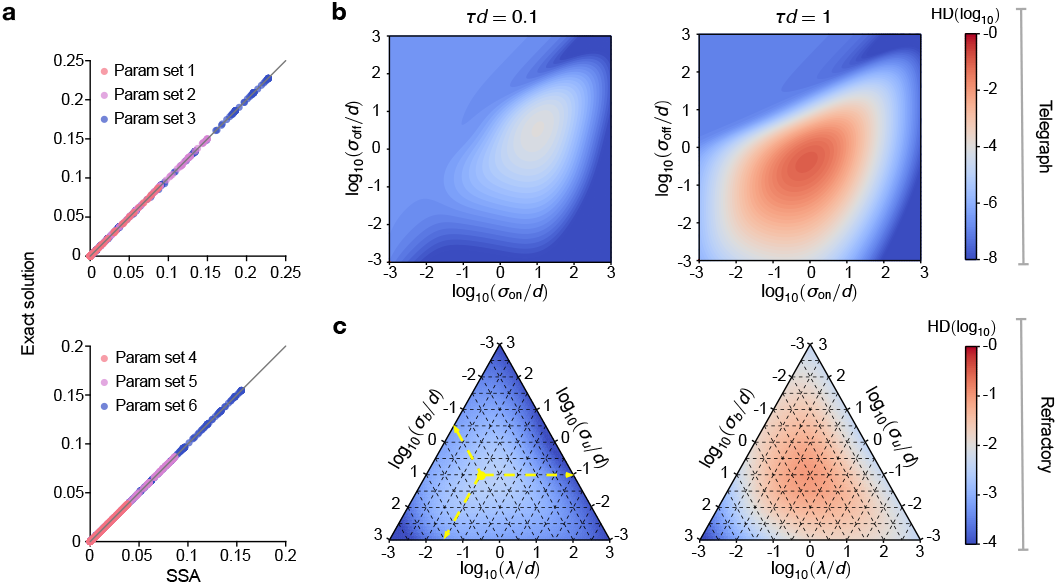
(a) P-P plots of the joint distributions computed from the solutions derived using queueing theory and the delay SSA. The exact solutions for Model I Eq. (9) (upper) are comprised of Eqs. (10) and (11), while those for Model II Eq. (12) (lower) are given by Eqs. (S55) and (11). The six sets of kinetic parameters are summarised in SM Table S1. (b) Hellinger distance (HD) between the joint distributions of the non-Markovian model (Model I) and its Markovian counterpart (Model III) as a function of the normalised gene state switching rates *σ*_on_*/d* and *σ*_off_*/d* (or *σ*_u_*/d, σ*_b_*/d* and *λ/d*), and the normalised export time *τd* (*ρ* is fixed to 5). (c) same as (b) but comparing Models II and IV. See SM Fig. S1 for the case *τd* = 10. Note that the yellow dashed arrows in (c) indicate how to read the ternary plot. In this figure the sum of the three logarithmic variables is equal to −3.

**Model II**. Another common stochastic transcriptional model is the refractory model, a three-state model in which burst initiation requires two steps [29]. It was developed to account for the experimental observation that for some mammalian genes the distribution of “off” intervals is not exponential but instead peaks at a nonzero value. By modifying it to a two-compartment version, the non-Markovian refractory model (referred to as Model II) consists of the following set of reactions

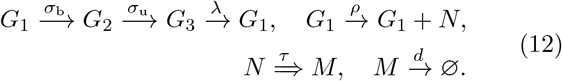

Applying the queueing theory result (Eq. (5)), we find the generating function solution of the stationary joint distribution is given by the sum of four _2_*F*_2_ hypergeometric functions (SM IV). Combining with Eq. (11), the joint distributions of mRNA numbers of Model II can be calculated, which is found to be in excellent agreement with the delay SSA (Fig. 2a lower panel).

### Comparison with Markovian models

Alternative competing models to the ones we solved are those where the export reaction 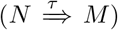 is modelled by a first-order reaction 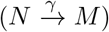, i.e. export time is an exponentially distributed random variable and hence the dynamics are Markovian [32, 42]. We will henceforth refer to the Markovian equivalent of Models I and II as Models III and IV, respectively. To meaningfully compare the two types of models, we set *γ* = 1/*τ* so that the mean nuclear export time is the same in both. In Fig. 2b we show the Hellinger distance between the steady-state joint distributions of Models I and III whereas in Fig. 2c we show the same but for Models II and IV. Note that the solution of the Markovian models is not known analytically and hence is computed numerically using finite state projection (FSP [43]). We find that the differences between the two types of models are very small when *dτ* is small (nuclear export is fast compared to cytoplasmic degradation). This is further substantiated by a theoretical analysis of the first two moments of mRNA fluctuations (SM V). An analysis of experimental data in Ref. [32] shows that about 80% of genes in mammalian cells have *dτ <* 1 (SM VI and SM Fig. S2) and hence suggests that in most cases, independent of the actual mechanism describing nuclear export, our queueing theory based solutions of Models I and II are accurate.

### A PGF-based parameter inference method

In principle, given the exact joint distribution of molecule numbers Eq. (11), one can use the method of maximum likelihood to estimate the transcriptional parameters from measurements of the nuclear and cytoplasmic mRNA in each cell. In practice this is not computationally efficient because this process involves a search over parameter space and for each point one needs to compute a large number of derivatives symbolically. To circumvent this issue, we develop an inference method that uses only the generating function solution. This consists of (i) constructing an empirical generating function from the data and (ii) finding the parameter set which minimizes the distance between the latter function and the analytical generating function. Given measurements of (*n*_*i*_, *m*_*i*_) counts of nuclear-cytoplasmic mRNAs for each cell in a population of *n*_*c*_ cells, it can be shown that the corresponding empirical generating function is given by

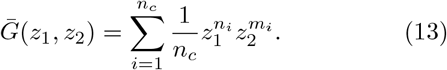

The inferred transcriptional parameters are then obtained by minimizing the following loss function:

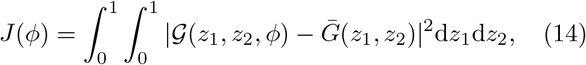

where 𝒢 (*z*_1_, *z*_2_, *ϕ*) is the analytical generating function of a user-defined gene expression model and 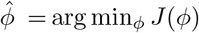 is the inferred parameter vector. Further details of the inference method are summarised in SM VII.

To understand the performance (including accuracy and computational efficiency) of the proposed inference method in the real experimental setup, we tested it on simulated data. Specifically, to closely mimic the experimental data collected from mammalian cells, we account for extrinsic noise due to the coupling of transcription to cell volume [44, 45] by choosing the transcription rate *ρ* to be proportional to the normalized cell volume *β* (cell volume divided by the average cell volume), i.e., the transcription reaction in the non-Markovian Model I reads 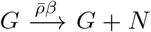. This implies that the burst size (*ρ/σ*_off_) increases with the cell volume, as has been shown by various experimental studies in eukaryotic cells [46, 47]. Furthermore, since eukaryotic cells have multiple copies of each gene, for simplicity we assume that each cell has two independent copies [48]. Our model is a simplification of more complex models of growing and dividing cells [49]. Given a parameter set and a volume distribution, one can then simulate Model I using the delay SSA [41] to generate *n*_*c*_ tuples of nuclear-cytoplasmic mRNA and cell volume observations (*n*_*i*_, *m*_*i*_, *β*_*i*_) which mimics experimental measurements from *n*_*c*_ cells. In Fig. 3a we show a cartoon that illustrates the setup. The model’s generating function solution (modified to account for extrinsic noise due to volume and two independent gene copies) is

**FIG. 3.**
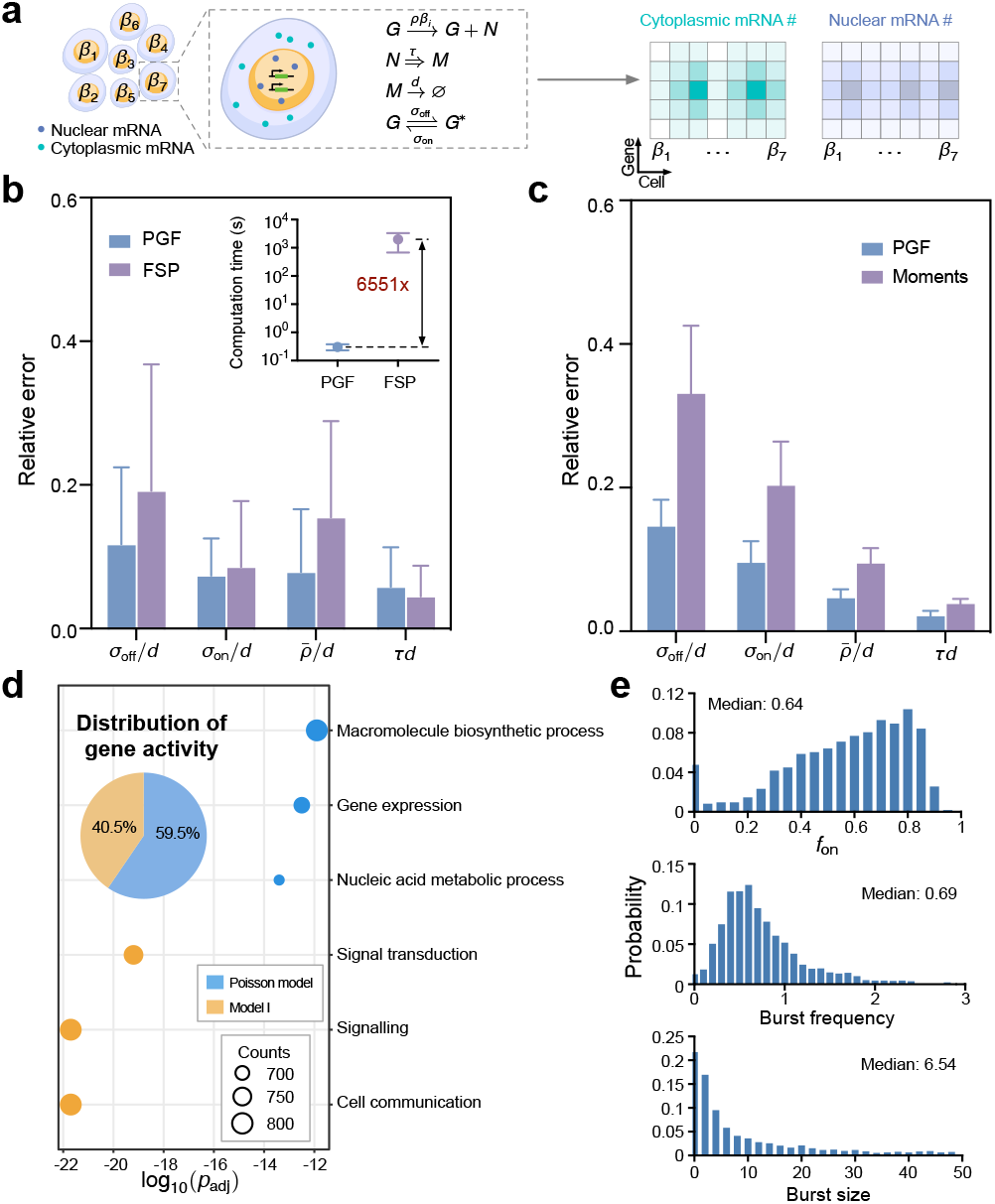
(a) Illustration of the simulated experimental setup used to test the PGF-based inference method. Each cell’s expression is described by an augmented version of Model I with two gene copies and a transcription rate that depends on the normalised cell volume *β* sampled from a gamma distribution. Delay SSA simulations generate joint count data for 15 batches of 2000 cells each with a different set of kinetic parameters. (b) Use of simulated data to compare the PGF-based inference method with maximum likelihood based inference using FSP shows comparable accuracy but higher computational efficiency: 0.30s (PGF) vs 1991.52s (FSP). The relative error is averaged over the 15 datasets and error bars show the standard error of the mean. (c) The PGF-based inference method is also more accurate than the method of moments. (d) Application of the PGF-based model selection method identifies the fraction of genes best fit by the Poisson model and Model I. This is followed by an ontology analysis which suggests the biological process functions associated with each of the two clusters of genes. (e) Application of the PGF-based inference method to fit the augmented version of Model I to nuclear and cytoplasmic MERFISH data for 2184 mammalian genes; these genes are better fitted by Model I than by the Poisson model and pass a fitting quality test on the moments (see Fig. S5). The inference output is shown by distributions of the fraction of time spent in the active state *f*_on_, the burst frequency (normalised by the degradation rate) and the burst size across the genome.

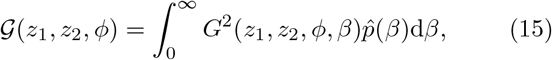

where the parameter vector is 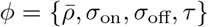 Note that the distribution of the normalized cell volume 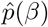 is estimated by fitting *β*_*i*_ with a kernel density estimation model (KDE) [50]. The inference method modified to account for extrinsic noise then consists of the optimization problem defined by Eq. (14) with 𝒢 (*z*_1_, *z*_2_, *ϕ*) given by Eq. (15). For implementation details see SM VIII A.

In Fig. 3b we show by the purple bars the relative errors of this inference method, given as input 15 different datasets obtained by simulation of Model I (SM Table S2) with the normalised cell volume distribution Gamma(7.5,0.11) [51]. The relative errors are small and comparable in size to those obtained by the conventional method of maximizing the likelihood (the joint probability distribution solution of the delay CME) numerically computed using the FSP method (SM VIII B) which are shown by the blue bars. However, the PGF method is remarkably faster, showing a speedup of several thousand times over FSP (inset of Fig. 3b) – this enhanced efficiency is particularly useful when fitting stochastic models to single-cell transcriptomic data for large numbers of genes, especially if the data is at sub-cellular resolution [52]. We also find that the PGF method is typically more accurate than another popular method based on matching the model’s first and second moments of mRNA fluctuations to those computed from data (Fig. 3c, SM VIII C and SM Table S3).

### Application of the PGF-based parameter inference method to experimental data

We consider single-cell nuclear and cytoplasmic mRNA count data for 6724 mammalian genes measured in 1368 human osteosarcoma cells [14] using multiplexed error-robust fluorescence in situ hybridization (MERFISH)(SM IX). Due to the absence of cell volume data, we used the total number of transcripts detected for each cell as a proxy for the cell volume (this is a reasonably good approximation as discussed in Ref. [53]); this is further divided by its average across the cell population to obtain the values of *β*_*i*_. Note that while we accounted for extrinsic noise due to volume, we did not account for technical noise; for this dataset the latter is minimal because transcript detection efficiency is high (80%) [14]. Since we did not *a priori* know if Model I is the best model to describe gene expression, we first fit the PGF of 2 competing models of gene expression to the empirical PGF of the data and then utilised 10-fold cross validation to determine the best fitting model (SM X). These 2 models are the Poisson model (constitutive expression; gene is always in a permissive state from which transcription can occur) and Model I (bursty expression; gene can switch from an off state to a permissive state from which transcription can occur). The former model is defined by the reaction schemes:

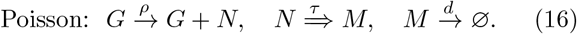

As before, extrinsic noise due to coupling of transcription to volume is introduced through *ρ* for Model I and the Poisson model (SM VIII). We found 4002 and 2722 genes that are best fit by the Poisson model and Model I, respectively. The corresponding fractions are summarised by the piechart in Fig. 3d inset. The high percentage of genes (about 60%) that are better fit by the Poisson model after controlling for extrinsic noise is in line with other recent studies [54–56]. Note that we did not fit Model II because experiments with simulated data showed that for a sample of size of few thousand cells it is not possible to easily tell apart Models I and II (Fig. S4). We further used the Database for Annotation, Visualization and Integrated Discovery (DAVID) [57] to identify biological functions related to the identified gene clusters. In Fig. 3d, we show that macromolecule production and signaling are associated with those genes best fit by the Poisson model and Model I, respectively. In Fig. 3e we show the inferred distributions of the burst size, burst frequency (normalised by the degradation rate) and the fraction of time spent in the active state *f*_on_ = *σ*_on_/(*σ*_on_ + *σ*_off_) of the genes that are best fit by Model I and which satisfy a reasonable fitting quality constraint (the average relative error across the first- and second-order moments of counts is less than 35%; see Fig. S5). The median *f*_on_ is 0.64, the median burst size is 6.83 and the median burst frequency (normalised by the degradation rate) is 0.69.

In SM Table S5 we repeat the model selection and parameter inference procedure for models with different assumptions and type of input data. Specifically we vary the number of gene copies assumed for Model I (4 instead of 2), vary the type of data (total counts per cell instead of joint count data) and the type of noise (intrinsic noise only instead of both intrinsic and extrinsic noise). We find that if we do not correct for extrinsic noise, the inference procedure leads to a large overestimation of the number of genes that are best fit by Model I (≈ 90% rather than ≈ 40% estimated earlier). For those genes which are best fit by Model I, the use of total count data rather than joint data leads to a significant underestimation of *f*_on_ and overestimation of the burst frequency. This makes a strong case for the use of transcriptomic data at sub-cellular resolution (joint data) for accurate parameter inference.

In summary, we have used queueing theory to derive an exact solution of the steady-state joint distribution of nuclear and cytoplasmic mRNA levels in a model of gene expression with general initiation mechanisms involving an arbitrary number of gene states. We have also devised a PGF-based parameter inference and model selection method that fits the exact PGF to the empirical PGF computed from transcriptomic data. Its application to a large experimental dataset led to a simple relationship between the type of gene expression and biological function.

## Acknowledgment

Z.C. acknowledges support from NSFC Grant (62073137), Shanghai Action Plan for Technological Innovation Grants (22ZR1415300, 22511104000, 23S41900500), the Natural Science and Engineering Research Council of Canada’s (NSERC’s) Discovery Grant (RGPIN-2024-06015) and Shanghai Center of Biomedicine Development. R.G. and J.S.-N. acknowledge support from a Leverhulme Trust grant (RPG-2020-327).

## Supplemental Material

### I. Derivation of the joint probability generating function for an arbitrary (general) interarrival time distribution

To compute *G*(*z*_1_, *z*_2_), we multiply *f*_*m*_(*x*) in Eq. (3) by 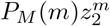 and sum it over all *m* ≥ 0, which gives

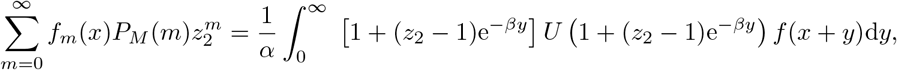

where *U* (*z*) is defined as

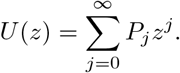

Using the following result derived in Ref. [37],

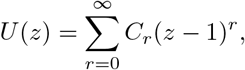

we get that

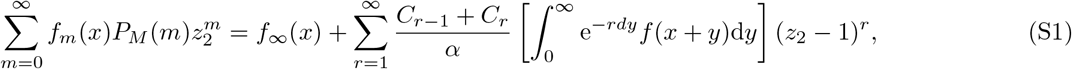

where *f*_∞_(*t*) is the probability density function of the forward recurrence time (the time until the next arrival measured from an arbitrary time point) in the stationary limit [30],

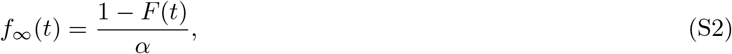

and *F* (*t*) is the cummulative probability function of the interarrival time, 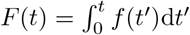. Next, we multiply 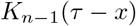 by 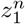 and sum it over all *n* ≥ 1, which gives

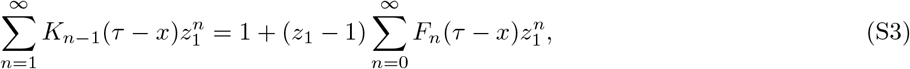

Where

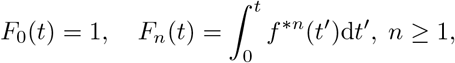

and *f* ^\**n*^ is the *n*-fold convolution of *f*. Let us denote by *P*_*N*_ (*n*) the probability distribution of nuclear RNA in the stationary limit. According to Ref. [38], *P*_*N*_ (*n*) is given by

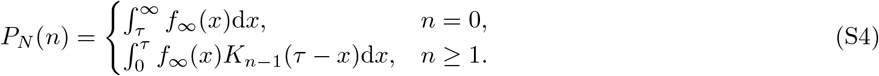

If we combine Eqs. (S1)-(S4), we get

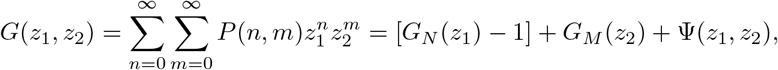

where *G*_*N*_ (*z*_1_) is the stationary probability generating function of nuclear mRNA number,

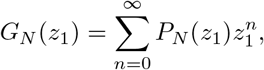

and Ψ(*z*_1_, *z*_2_) is defined as

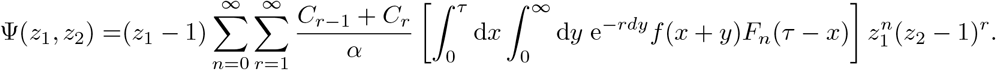

Let us denote by *ϕ*(*rd, x*) the following integral,

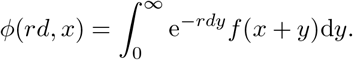

The expression for Ψ(*z*_1_, *z*_2_) can now be written more compactly as

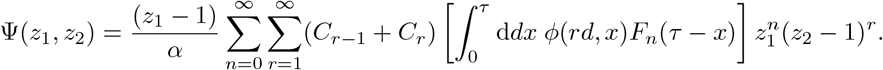

By taking the Laplace transform of Ψ(*z*_1_, *z*_2_) with respect to *τ* we get

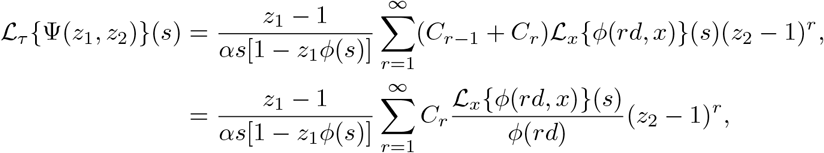

where we used that *C*_*r*−1_ + *C*_*r*_ = *C*_*r*_/*ϕ*(*rd*) [37] and ℒ_*x*_ *ϕ*(*rd, x*) (*s*) is the Laplace transform of *ϕ*(*rd, x*) with respect to *x*,

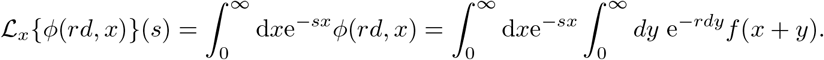

### II. Derivation of Eq. (10) using queueing theory

The probability density of the time between two nuclear mRNA production events for the reaction system Eq. (9) is a phase-type distribution of the form

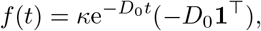

with *κ* = [0, 1]^⊤^, **1** = [1, 1]^⊤^ and

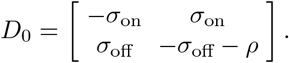

The corresponding Laplace transform of *f* (*t*) is

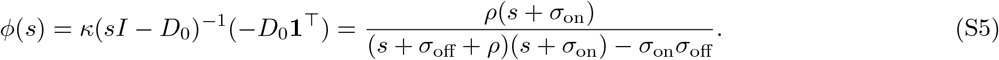

The mean interarrival time *α* follows from *α* = −*ϕ*′(0) and hence is given by

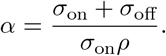

By taking the inverse Laplace transform of Eq. (7), we obtain the probability generating function of nuclear Mrna

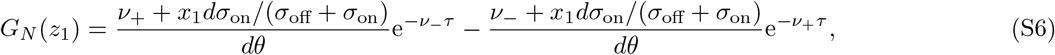

with 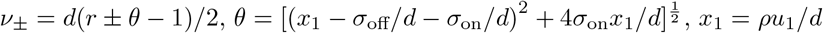 and *u*_1_ = *z*_1_ − 1.

It follows from Eqs. (4) and (S5) that

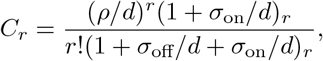

which together with Eq. (6) leads to the probability generating function of cytoplasmic mRNA

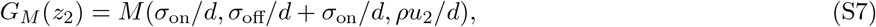

where *u*_2_ = *z*_2_ − 1 and *M* is Kummer’s confluent hypergeometric function.

To calculate *G*(*z*_1_, *z*_2_) using Eq. (5), we need to calculate Ψ(*z*_1_, *z*_2_). Using

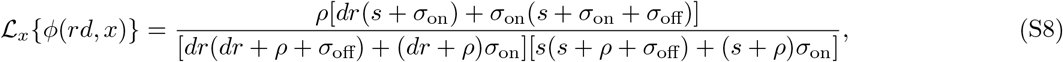

we find that the Laplace transform of Ψ(*z*_1_, *z*_2_) (as given by Eq. (8)) is given by

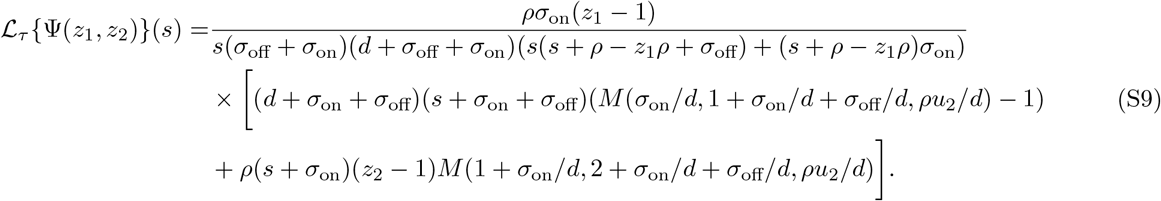

Substituting the inverse Laplace transform of Eq. (S9), and Eqs. (S6) and (S7) in Eq. (5), we find after simplification using `Mathematica` [63] that the solution for *G*(*z*_1_, *z*_2_) is given by Eq. (10).

### III. Derivation of Eq. (10) using the delay master equation

#### A. Derivation of the delay master equations

Consider the system of reactions given by Eq. (9). To derive the delay master equations describing the stochastic dynamics, we want to obtain the probability *P* (*φ, n, m, t* + Δ*t*), i.e., the probability of observing *n* nuclear mRNAs and *m* cytoplasmic mRNAs at time *t* + Δ*t* given that the gene state is *φ*. Note that the inactive and active gene states are denoted by *φ* = 0 and *φ* = 1, respectively. There are four sets of events which contribute to this probability in the time interval (*t, t* + Δ*t*):

A. a nuclear mRNA transcript is produced while the gene is active;
B. either the gene state changes or a cytoplasmic mRNA is degraded;
C. a nuclear mRNA transcript is exported to the cytoplasm and becomes a cytoplasmic mRNA;
D. no reaction occurs.

In the infinitesimal limit Δ*t* → 0, the probability of event (A) simply follows the conventional propensity associated with a firing of the transcription reaction:

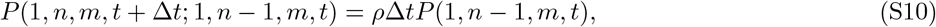

where *P* (*φ, n, m, t*; *φ*′, *n*′, *m*′, *t*′) is the joint probability of having gene state *φ, n* nuclear mRNAs and *m* cytoplasmic mRNAs at time *t* and gene state *φ*′, *n*′ nuclear mRNAs and *m*′ cytoplasmic mRNAs at time *t*′. Similarly, the joint probabilities of the reactions associated with event (B) are

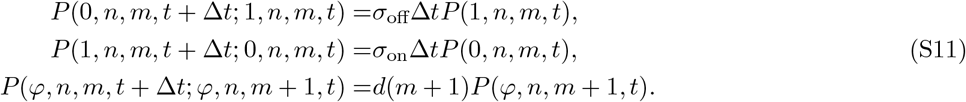

The probability of event (C) occurring requires a careful consideration of the history of the process: (i) (1, *n*′, *m*′, *t*) → (1, *n*′ + 1, *m*′, *t* − *τ* + Δ*t*); (ii) (1, *n*′ + 1, *m*′, *t* − *τ* + Δ*t*) → (*φ, n* + 1, *m* − 1, *t*); (iii) (*φ, n* + 1, *m* − 1, *t*) → (*φ, n, m, t* + Δ*t*), where *n*′ and *m*′ are the number of nuclear and cytoplasmic RNAs at time *t* − *τ*. The elementary probability of event (C) can hence be decomposed into

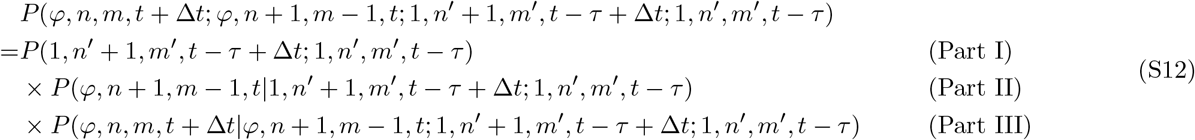

The probability of Part III is 1 since every production event is followed by an export event a time *τ* later, i.e.,

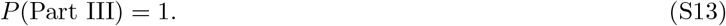

According to Eq. (S10), the probability of Part I is

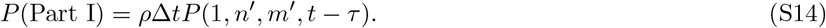

The two “+1”s of nuclear mRNAs in the probability of Part II represent a nuclear mRNA was born in time (*t* −*τ, t*− *τ* + Δ*t*) and stays in the system till time *t*, and notably does not participate in any reaction in time (*t* − *τ* + Δ*t, t*). Thus, we have

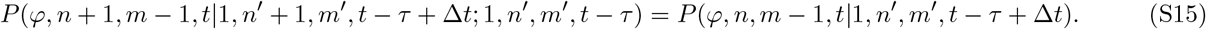

Since all nuclear mRNAs born before time *t* − *τ* (of which the number is *n*′) are exported from nucleus to cytoplasm, the number of nuclear mRNAs (*n*) at time *t* is independent of *n*′ and *m*′, but only depends on the gene state at time *t* − *τ*. Besides, the number of cytoplasmic mRNAs (*m* − 1) at time *t* is determined by the number of nuclear mRNAs out of *n*′ exported to cytoplasm and the number of cytoplasm RNA out of *m*′ degraded in the time interval (*t* − *τ* +Δ*t, t*). As such, we have

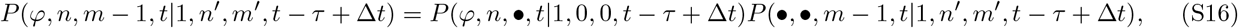

where the symbol • stands for the marginalization over the variable in the same position. For instance,

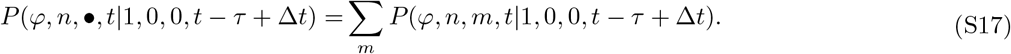

It follows that by means Eqs. (S12)-(S17) that the total probability of event (C) is given by

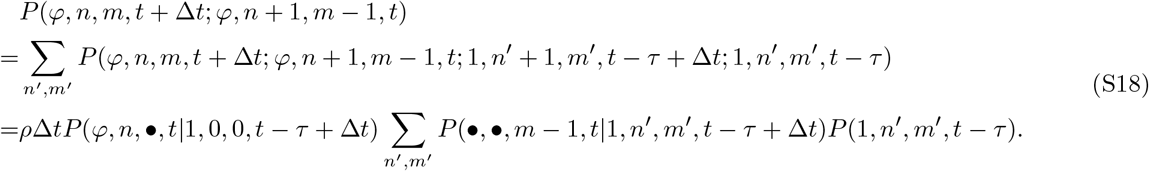

Note that here we used the law of total probability. See Fig. S3 for an illustration of the history of event (C). The probability that event (D) occurs is obtained by subtracting all the probabilities of one reaction occurring from *P* (*φ, n, m, t* + Δ*t*). Specifically, we have

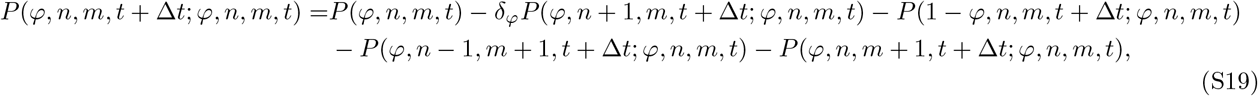

where *δ*_*φ*_ is Kronecker delta, and *δ*_*φ*_ = 1 if and only if *φ* = 1.

By means of Eqs. (S10), (S11), (S18) and (S19), it can be shown that

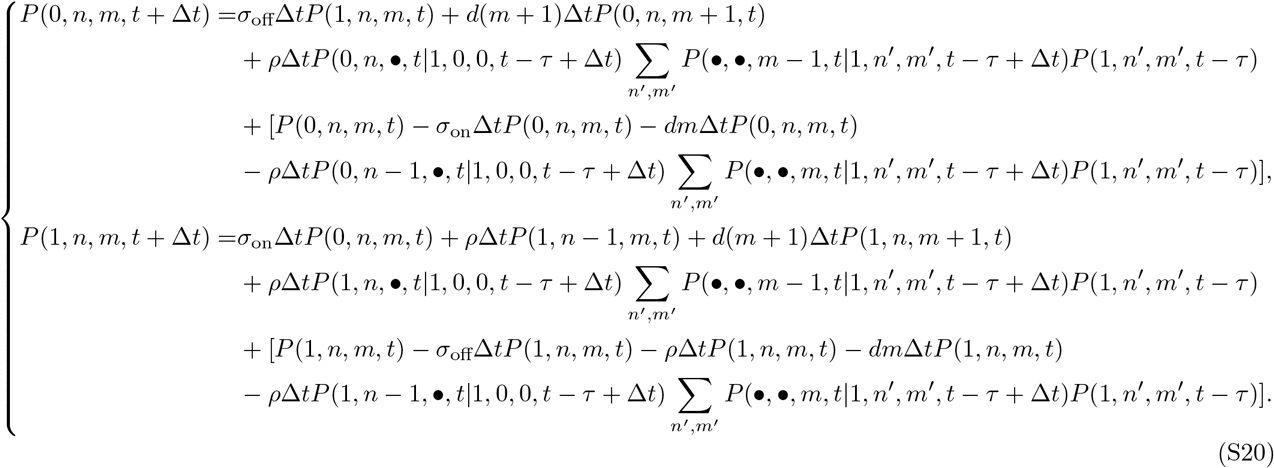

Dividing both sides of Eq. (S20) by Δ*t* and letting Δ*t* →0, the delay master equation of the delayed RNA export model is obtained:

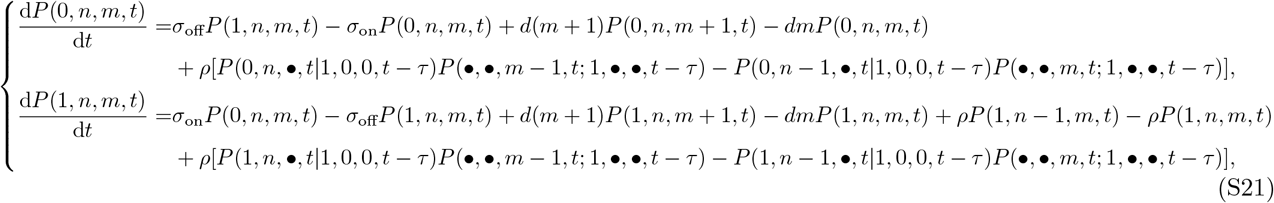

in which the following simplification step is used

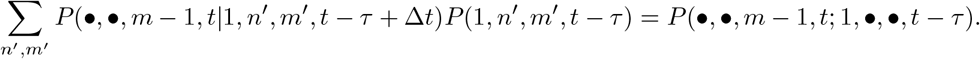

Next we seek solutions to the two probabilities, i.e., *P* (*φ, n*, •, *t*|1, 0, 0, *t* − *τ*) and *P* (•, •, *m* − 1, *t*; 1, •, •, *t* − *τ*). The *n* nuclear mRNAs of the two conditional probabilities *P* (*φ, n*, •, *t*|1, 0, 0, *t* − *τ*) for *φ* = 0, 1 in Eq. (S21) are produced during time (*t* − *τ, t*), and the pertinent dynamics are that of a reaction system composed of the first three reactions in Eq. (9), i.e., *G* ⇌ *G*^*^ and *G* → *G* + *N*. Specifically, we have

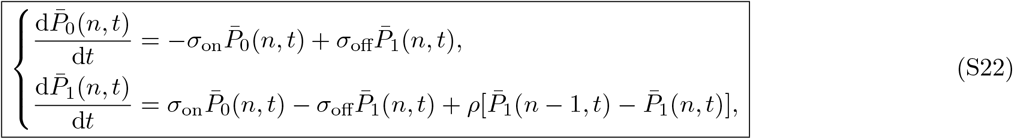

with initial conditions 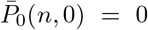 for any 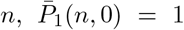 for *n* = 0 and equal to 0 otherwise, as well as 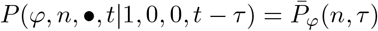 for *φ* = 0, 1.

By summing over all possible *n* on both sides of Eq. (S21), we obtain

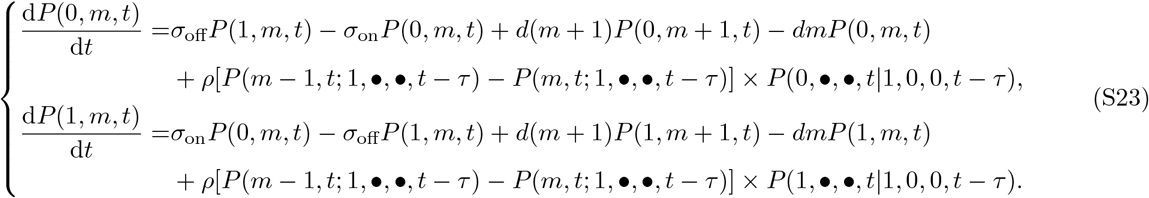

Summing up the two equations in Eq. (S23) leads to

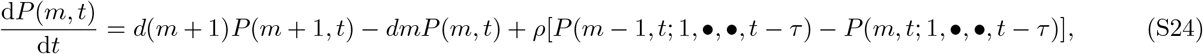

where *P* (*m, t*) = Σ_*φ*_ *P*_*φ*_(*m, t*). Given the nature of the delayed export, the dynamics of cytoplasmic mRNAs is equivalent to that of a conventional telegraph model a *τ* time earlier. The telegraph model encompasses the following set of reactions

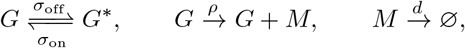

of which the governing master equations are

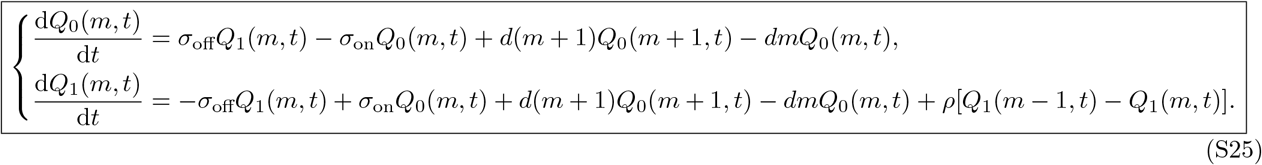

Also by summing up the two equations in Eq. (S25), we have that

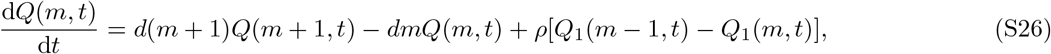

where *Q*(*m, t*) =Σ _*φ*_ *Q*_*φ*_(*m, t*). Noted that *P* (*m, t*) = *Q*(*m, t − τ*), and by comparing Eqs. (S24) and (S26), we conclude that for *t > τ*

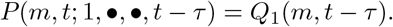

By replacing the two conditional probabilities in Eq. (S21) with 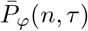 and *Q*_1_(*m, t* − *τ*), Eq. (S21) becomes

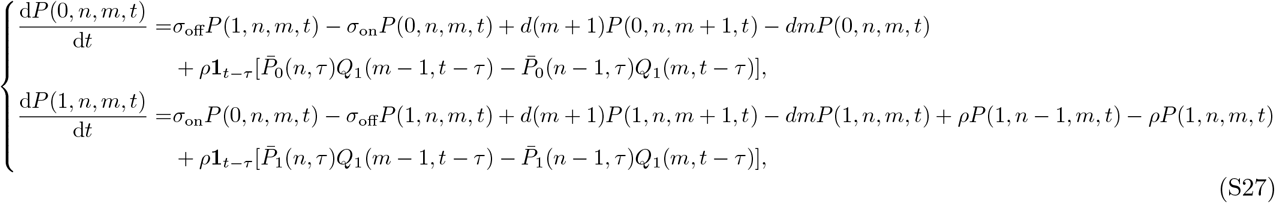

which together with Eqs. (S22) and (S25) fully characterizes the stochastic dynamics of the system of reactions in Eq. (9). Note that in Eq. (S27), **1**_*t*−*τ*_ = 1 if and only if *t > τ*. Note that the delay master equation [64, 65] is not the standard master equation [66], because the presence of a deterministic export time means that the model is non-Markovian.

#### B. Solution conditional on the gene state

In this subsection, we solve Eq. (S27) in steady state conditions by means of the generating function method. By defining generating functions 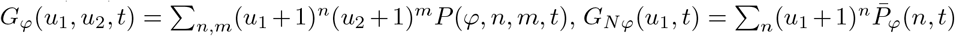 and *G*_*Mφ*_(*u*_2_, *t*) = Σ _*m*,_ (*u*_2_ + 1)^*m*^*Q*_*φ*_(*m, t*), Eqs. (S22), (S25) and (S27) are transformed into

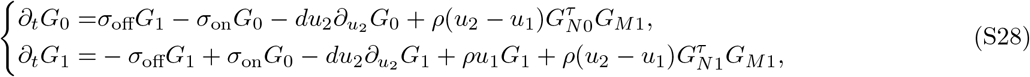

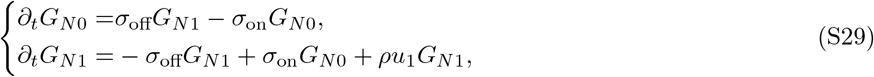

And

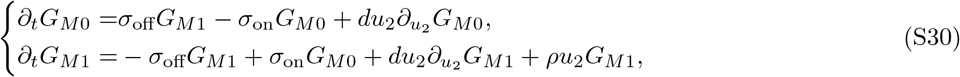

where the arguments *u*_1_, *u*_2_ and *t* are suppressed for brevity, and 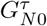 is short for *G*_*N*0_(*u*_1_, *τ*). Prior to solving Eq. (S28), we need to find the solutions to Eqs. (S29) and (S30).

The exact solution to Eq. (S30) has been reported in Ref. [67]. At steady state, Eq. (S30) admits the solution

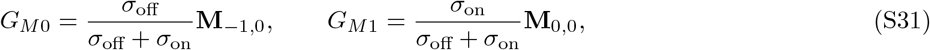

where *x*_2_ = *ρu*_2_/*d* and the Kummer function

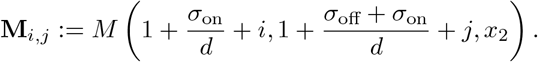

Note that Eq. (S29) consists of two ordinary differential equations (ODEs). We solve *G*_*N*1_ as a function of *G*_*N*0_ from the first equation in Eq. (S29) and insert the solution into the second one to obtain a second-order ODE. Using initial conditions *G*_*N*0_ = 0 and *G*_*N*1_ = 1 at time *t* = 0 we then solve the second-order ODE at time *τ* to obtain

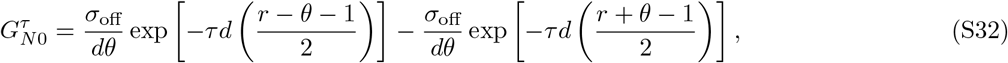

with *x*_1_ = *ρu*_1_/*d, α* = *σ*_on_/(*σ*_off_ + *σ*_on_), *r* = 1 + (*σ*_off_ + *σ*_on_)/*d* − *x*_1_ and

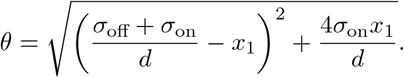

The probability associated with the active gene state immediately follows

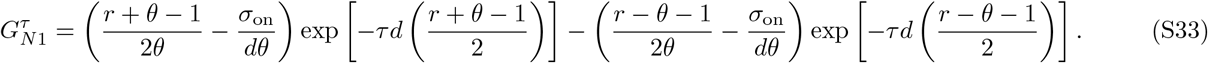

Letting ∂_*t*_*G*_0_ = ∂_*t*_*G*_1_ = 0, we solve *G*_1_ as a function of *G*_0_ from the first equation in Eq. (S28), and have that

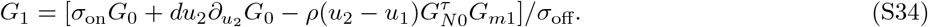

Summing up the two equations in Eq. (S28) leads to

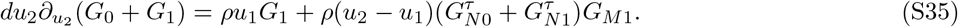

Inserting Eqs. (S31) and (S34) into Eq. (S35), we then get an equation that only depends on *G*_0_:

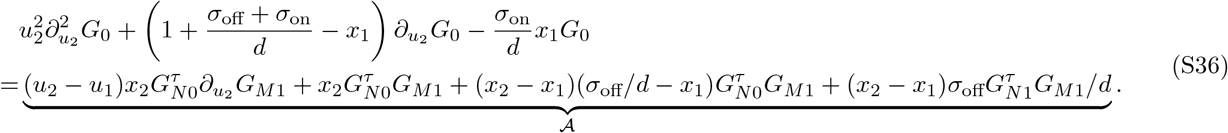

By means of Eqs. (13.3.3), (13.3.4) and (13.3.15) in Ref [68], the term A simplifies to

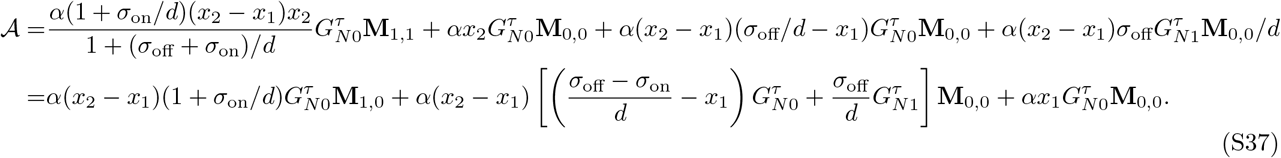

It can be ascertained that Eq. (S36) is a linear inhomogeneous second-order ODE. Therefore, the solution to Eq. (S36) is a combination of solutions to the general ODEs of the following form

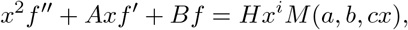

where *A, B, H, a, b, c* are constants and *i* = 0, 1. The general solution is given by

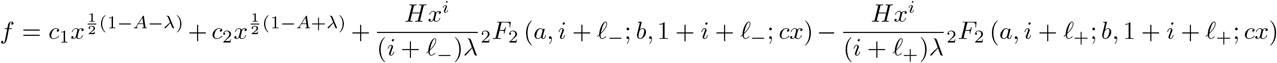

with 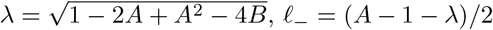 and 𝓁_+_ = (*A* − 1 + *λ*)/2. By defining the hypergeometric function

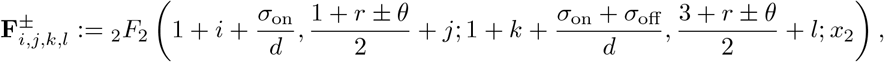

the general solution to Eq. (S36) reads

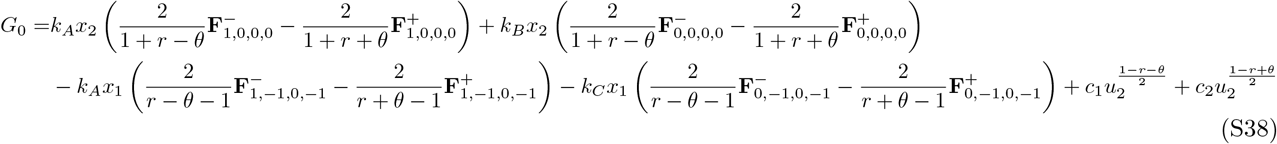

Where

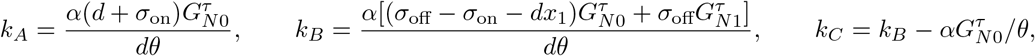

and constants *c*_1_ and *c*_2_ remain to be determined.

Since 1 − *r* − *θ <* 0 and *u*_2_ ∈ [−1, 0], the term 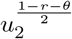 becomes ill-defined when *u*_2_ approaches 0 from the negative real axis, thereby indicating *c*_1_ = 0. Additionally, (1 − *r* + *θ*)/2 can hardly be an integer, and the generating function is a polynomial of only integer powers, thereby suggesting *c*_2_ = 0.

By plugging Eq. (S38) into Eq. (S34), we obtain an equation for *G*_1_ of the form

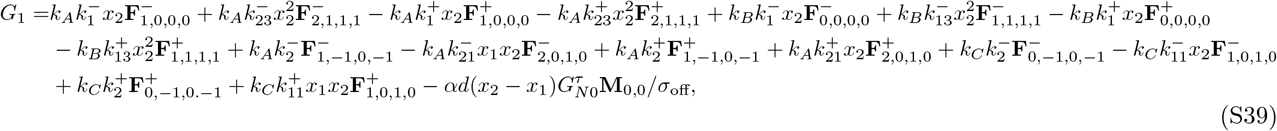

With

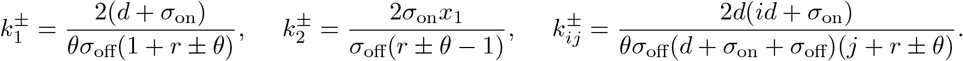

#### C. Solution independent of the gene state

Since the solutions Eqs. (S38) and (S39) are rather complicated, we derive much simpler solutions that are independent of the gene state. Note that the solutions Eqs. (S32) and (S33) can be compactly written as

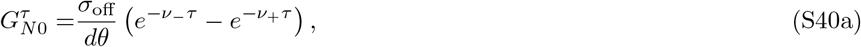

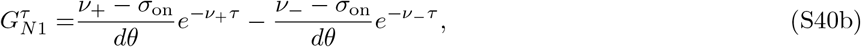

using the definition *ν*_±_ = *d*(*r* ± *θ* −1)/2.

Additionally, by combining the two equations in Eq. (S28) in the steady-state limit, we obtain the following ODE for the joint probability generating function *G* = *G*_0_ + *G*_1_,

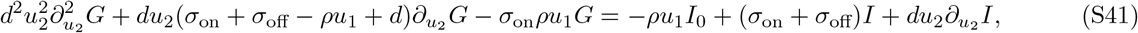

where 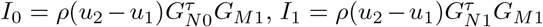 and *I* = *I*_0_ + *I*_1_. The solution to this equation takes a simple form in terms of two Kummer functions, which precisely agrees with Eq. (10), a result that has been previously derived using queueing theory.

### IV. Stochastic description of reaction system Eq. (12) using queueing theory

Without loss of any generality, here we present the solution for *d* = 1. The probability density of the time between two successive nuclear mRNA production events for the reaction system Eq. (9) is a phase-type distribution,

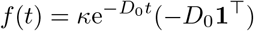

with *κ* = [0, 0, 1]^⊤^, **1** = [1, 1, 1]^⊤^ and

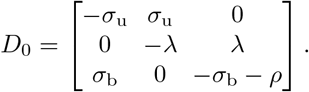

The corresponding Laplace transform of *f* (*t*) is

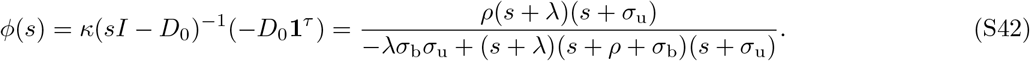

The mean interarrival time *α* follows from *α* = −*ϕ*′(0) and is given by

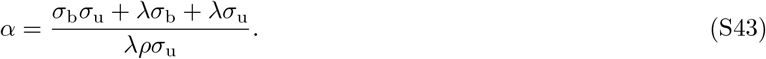

Using Eqs. (7), (S42) and (S43), the Laplace transform of *G*_*N*_ (*z*_1_) with respect to the nuclear processing time *τ* is

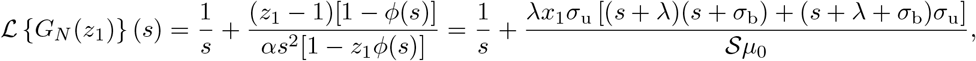

where *u*_1_ = *z*_1_ − 1, *x*_1_ = *ρu*_1_, *µ*_0_ = *σ*_b_*σ*_u_ + *λσ*_b_ + *λσ*_u_ and

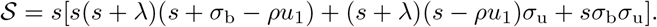

S = *s*[*s*(*s* + *λ*)(*s* + *σ*_b_ − *ρu*_1_) + (*s* + *λ*)(*s* − *ρu*_1_)*σ*_u_ + *sσ*_b_*σ*_u_].

It follows from Eq. (4) that *C*_*j*_ for *i* = 1, 2, …are given by

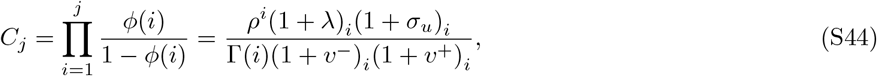

Where

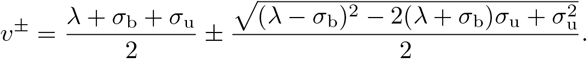

Using Eqs. (6) and (S44), we obtain the probability generating function for cytoplasmic Mrna

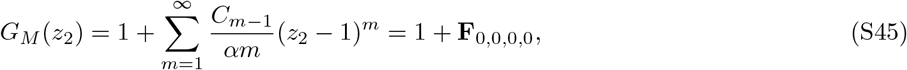

where *u*_2_ = *z*_2_ − 1, *x*_2_ = *ρu*_2_ and **F**_*i,j,k,l*_ = _2_*F*_2_ (*λ* + *i, σ*_u_ + *j*; *v*^−^ + *k, v*^+^ + *l*; *x*_2_). Next we compute the Laplace transform ℒ {*ϕ*(*r, x*)} as

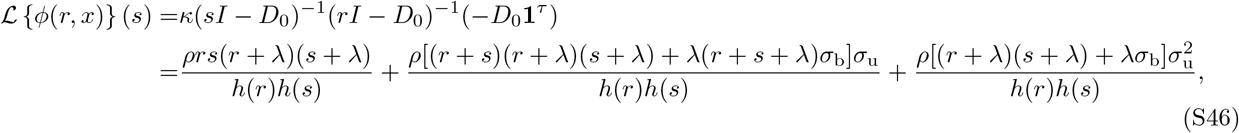

where *h*(*x*) = *x*(*x* + *λ*)(*x* + *ρ* + *σ*_b_) +(*x* + *λ*)(*x* + *ρ*)*σ*_u_ + *xσ*_b_*σ*_u_. According to Eq. (S46), the summation term in Eq. (8) becomes

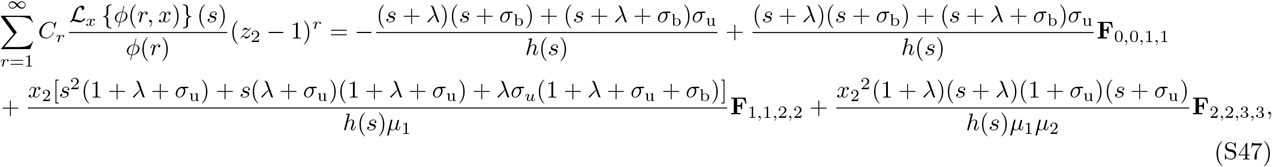

where *µ*_1_ = (1 + *σ*_b_)(1 + *σ*_u_) + *λ*(1 + *σ*_b_ + *σ*_u_), *µ*_2_ = (2 + *σ*_b_)(2 + *σ*_u_) + *λ*(2 + *σ*_b_ + *σ*_u_). Given Eq. (S47), the Laplace transform of Ψ(*z*_1_, *z*_2_) becomes

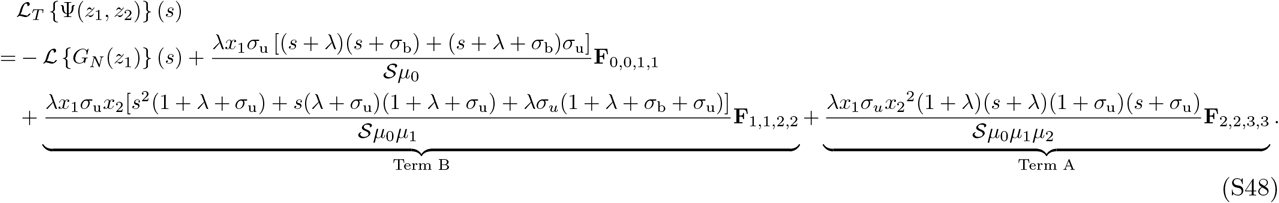

Next we simplify Terms A and B using the relation *bz*_2_*F*_2_(*a* + 1, *b* + 1; *c* + 1, *d* + 1; *z*) = *cd*_2_*F*_2_(*a* + 1, *b*; *c, d*; *z*) − *cd*_2_*F*_2_(*a, b*; *c, d*; *z*) [69]. This leads to

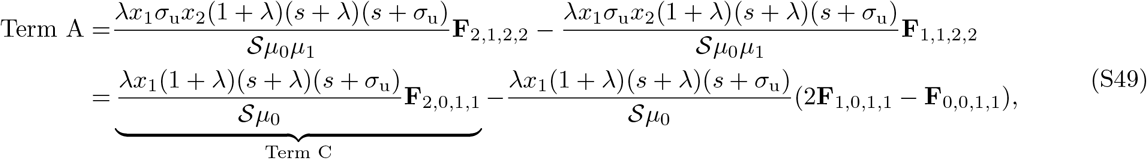

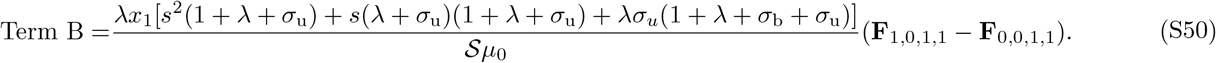

Given another relation, *a*_2_*F*_2_(*a* + 1, *b*; *c, d*; *z*) = *b*_2_*F*_2_(*a, b* + 1; *c, d*; *z*) + (*a* − *b*)_2_*F*_2_(*a, b*; *c, d*; *z*) [69], we simplify Term C to

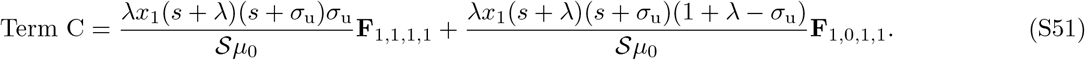

Using Eqs. (S49), (S50) and (S51), Eq. (S48) can be simplified to

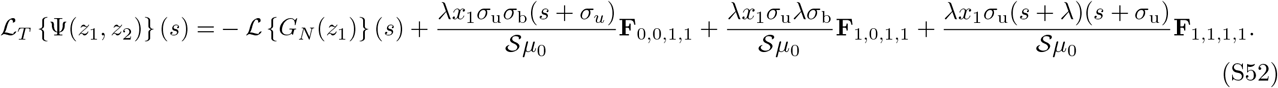

Subsequently, we need to calculate the inverse Laplace transform of ℒ_*T*_ {Ψ(*z*_1_, *z*_2_)} (*s*). Note that the last three terms in Eq. (S52) share the same denominator, of which the poles are denoted here by *r*_1_, *r*_2_, *r*_3_; these are the roots of the equation *S* = 0. By using the residue method, we derive the following results:

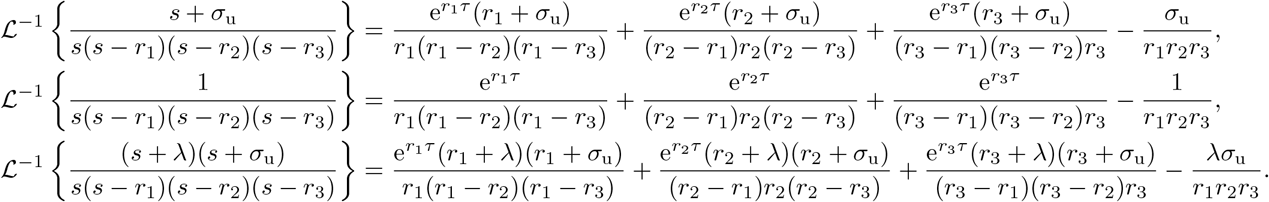

Using these equations, the function Ψ(*z*_1_, *z*_2_) is found to be

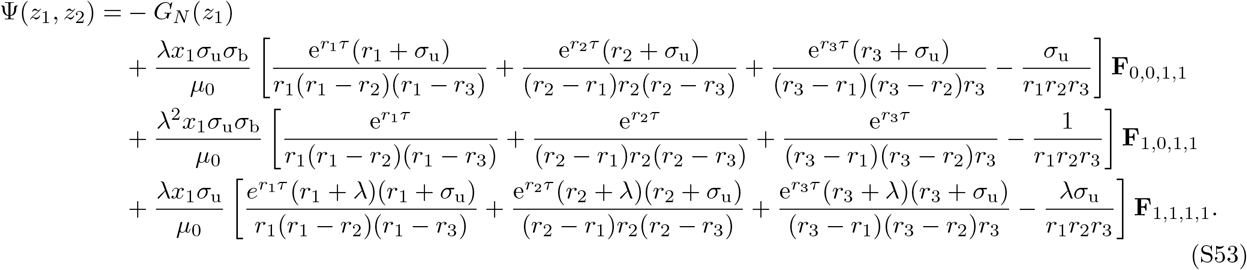

Solving the three roots *r*_1_, *r*_2_ and *r*_3_ of the equation 𝒮 = 0 we find

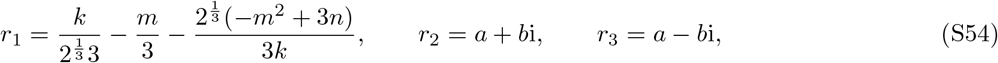

Where

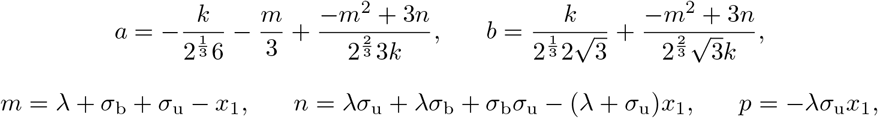

and

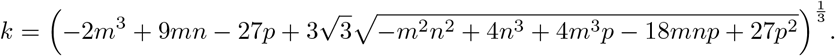

Finally using Eqs. (S53), (S54) and (5), the generating function solution is found to be given by

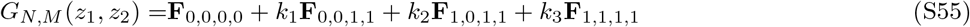

together with

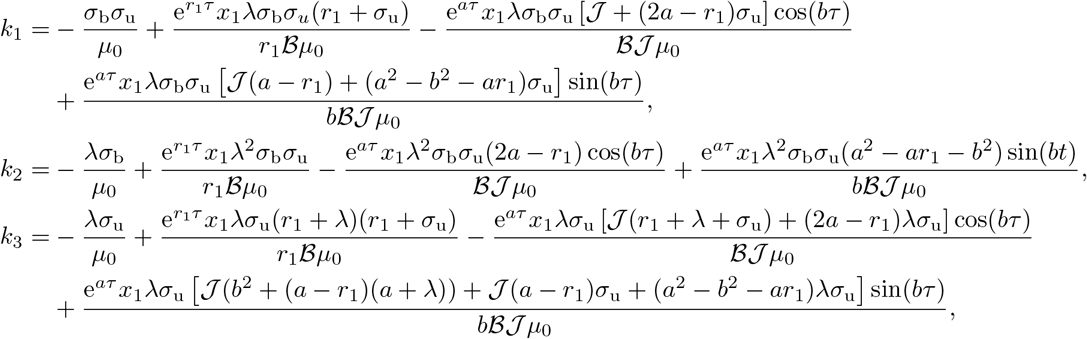

and 𝒥= *a*^2^ + *b*^2^, B = (*a* − *r*_1_)^2^ + *b*^2^.

### V. The first and second-order moments of Models I and III converge under fast export conditions

By taking the first-order derivatives of Eq. (10) with respect to *z*_1_ and *z*_2_ and letting *z*_1_ = 1 and *z*_2_ = 1, we obtain the first-order moments of Model I:

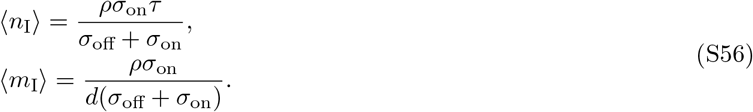

From the master equation of Model III (or the rate equations) we can obtain the first-order moments, which are

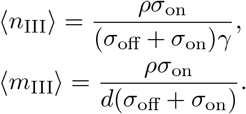

Since *γ* = 1/*τ*, it immediately follows that that ⟨*n*_I_⟩ = ⟨*n*_III_⟩ and ⟨*m*_I_⟩ = ⟨*m*_III_⟩.

Similarly, by taking the second-order derivatives of Eq. (10) with respect to *z*_1_ and *z*_2_ and letting *z*_1_ = 1 and *z*_2_ = 1, all the second-order moments of Model I can be obtained

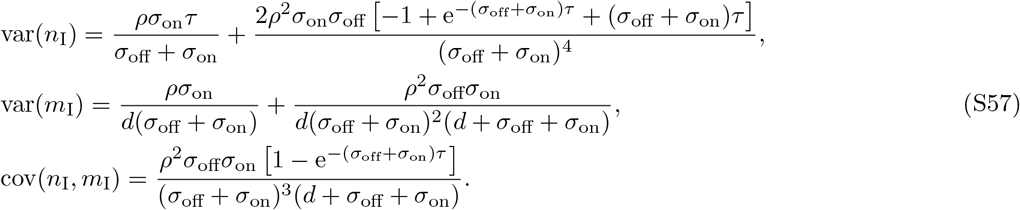

Using the master equation of Model III we can also derive the exact second-order moments, which are

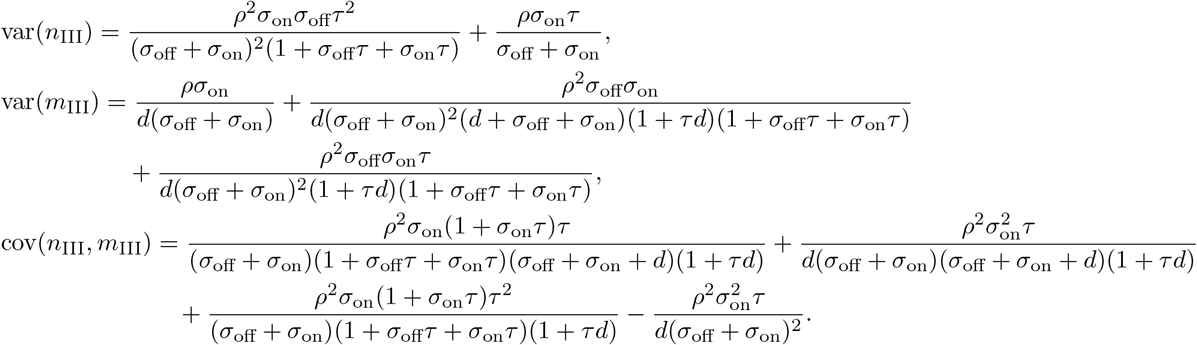

By expanding all the second-order moments of Models I and III as a series in powers of *τ* about *τ* = 0, we find that

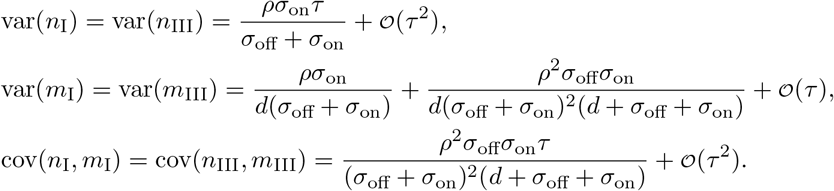

Hence given only information about the first and second moments of mRNA counts, the two models become indistinguishable in the limit of fast nuclear export.

### VI. Estimation of the ratio of export to degradation times from experimental data

The steady-state means of nuclear and cytoplasmic mRNA counts according to Model I are given by Eq. (S56). This implies that ⟨*n*⟩ / ⟨*m*⟩ = *τd*. To estimate *τd*, we extracted the average numbers of nuclear and cytoplasmic mRNAs in mouse MIN6 cells and Liver cells from Table S2 of Ref. [32]. We found that approximately 80% of genes for which the mean number of nuclear and cytoplasmic mRNAs is not zero, are characterised by *τd <* 1 (Fig. S2).

### VII. Parameter inference using the generating function solution for synthetic data without extrinsic noise and for one gene copy

Consider a sample of *n*_*c*_ cells in steady-state conditions. Specifically, in each cell *i*, we measure its nuclear mRNA counts (*n*_*i*_) and cytoplasmic mRNA counts (*m*_*i*_). The empirical joint distribution for the cell population is

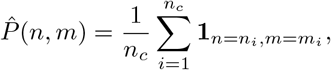

where 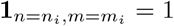 if and only if *n* = *n*_*i*_ and *m* = *m*_*i*_. The empirical generating function for the cell population is then computed as

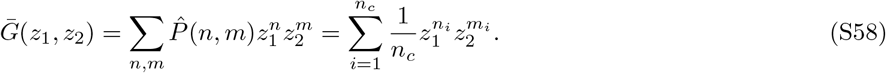

To infer the kinetic parameters *ϕ*, we aim to minimize the loss function in the form of the *L*^2^ norm

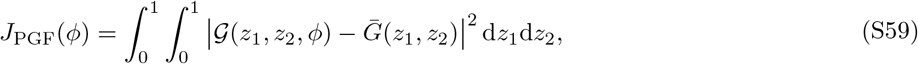

where 𝒢 (*z*_1_, *z*_2_, *ϕ*) is the generating function of the model. For the Model I (Eq. (9)), 𝒢 (*z*_1_, *z*_2_, *ϕ*) is the generating function solution given by Eq. (10), and *ϕ* = {*ρ, σ*_on_, *σ*_off_, *τ*}; for the Poisson model Eq. (16),

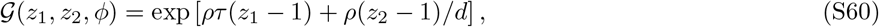

and *ϕ* = {*ρ, τ*}. Note that since we are only considering the steady state limit, only the kinetic parameters normalised by *d* can be estimated. Note also that Eq. (S60) can be derived from the general queueing theory result, Eq. (5).

To improve computational efficiency, we utilize the Gauss quadrature method and approximate the integral by

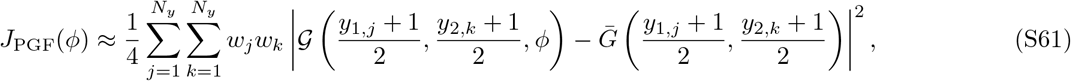

where *y*_1,*j*_ ∈ [−1, 1] and *y*_2,*k*_ ∈ [−1, 1] are the *j*-th and *k*-th integral points of the *N*_*y*_-th order Gauss quadrature approximation, and *w*_*j*_, *w*_*k*_ are the corresponding integral weights given by the command `gausslegendre` in Julia.

Finally, we employ the Nelder-Mead algorithm to minimize *J*_PGF_(*ϕ*). We use this algorithm because: (i) it is more robust against the selection of initial conditions, particularly compared to gradient-based optimization algorithms; (ii) the computation of the gradients ∂_*ϕ*_*J*_PGF_(*ϕ*) is rather time demanding. The optimization is implemented in the package `Optim.jl` in Julia.

The numerical recipe of the generating-function-based inference method is summarised in Algorithm 1. In this study, the default threshold used in the Julia command `optimize` is gradient tolerance `g_tol`= 10^−20^ (the absolute value of the gradient of the loss function *J*_PGF_(*ϕ*) less than 10^−20^) and maximal iterations `iterations`= 1000.

#### Algorithm 1

Generating-function-based parameter inference for nuclear-cytoplasmic mRNA count data

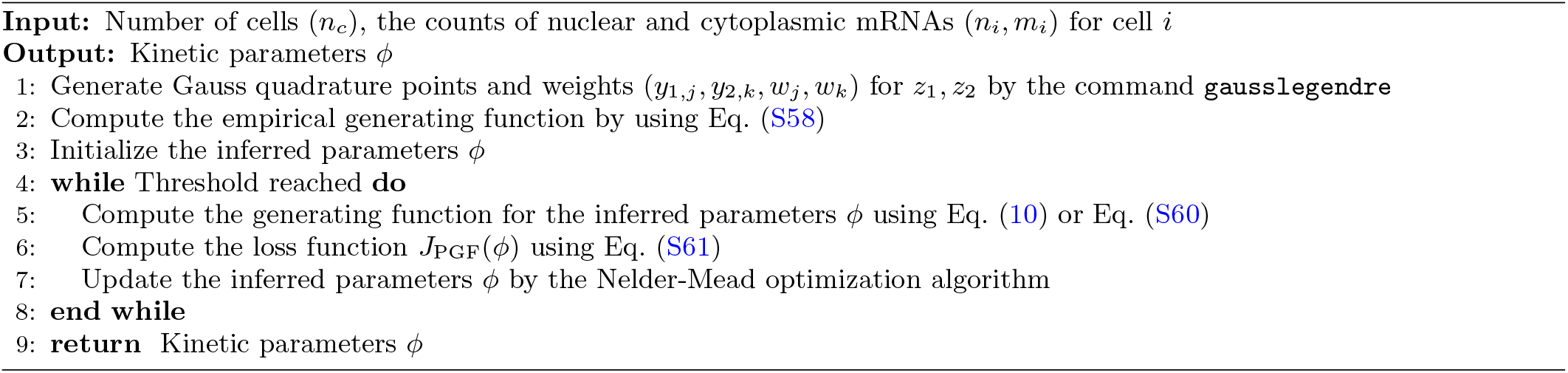

### VIII. Parameter inference for data with extrinsic noise and multiple gene copies

Consider a sample of *n*_*c*_ cells in steady-state conditions. Specifically, in each cell *i*, we measure its nuclear mRNA counts (*n*_*i*_), cytoplasmic mRNA counts (*m*_*i*_) and cellular volume (*V*_*i*_). The empirical distribution for the cell population is calculated using Eq. (S58). The normalized volume for cell *i* is given by

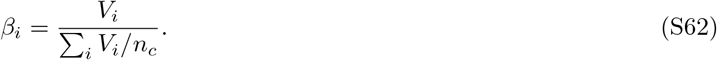

Because of extrinsic noise due to the coupling of the transcription rate to cell volume, a common property of many eukaryotic cells [46, 47], the transcription step in Model I (Eq. (9)) and the Poisson model (Eq. (16)) is modified as

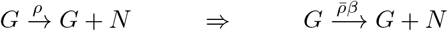

where *β* is a random variable sampled from an unknown distribution *p*(*β*), representing the transcriptional extrinsic noise. The unknown distribution is estimated by using the kernel density estimation (KDE) method [50] and the estimate is denoted as 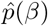. For Model I (Eq. (9)), the kinetic parameters 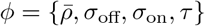 are the ones which we will infer; for the Poisson model (Eq. (16)), 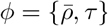.

#### A. Generating-function-based parameter inference

Let the joint distribution of nuclear and cytoplasmic mRNA counts (*n, m*) for cell *i* according to a model be *P* (*n, m, ϕ, β*_*i*_), and let the corresponding generating function for a single copy gene in cell *i* be *G*(*z*_1_, *z*_2_, *ϕ, β*_*i*_). Assuming each cell has two independent copies of each gene, the generating function used in Eq. (S59) for the cell population reads

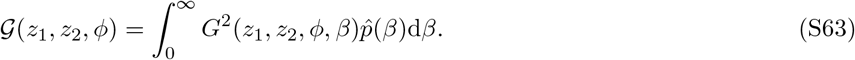

As before, to increase computational efficiency we utilize the Gauss quadrature method and transform the integral variable, thereby Eq. (S63) becomes

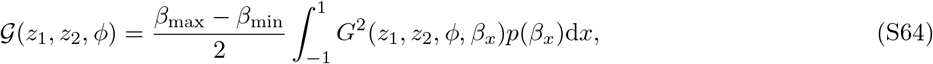

Where

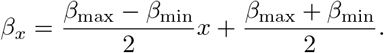

Here *β*_max_ = max_*i*_ *β*_*i*_, *β*_min_ = min_*i*_ *β*_*i*_ and *x* is the new integral variable. The numerical approximation to Eq. (S64) is then

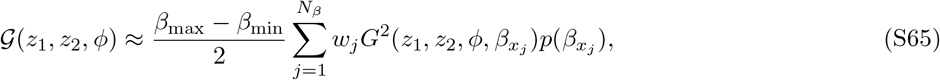

where *x*_*j*_ is the *j*-th integral point of the *N*_*β*_-th order Gauss quadrature approximation, and *w*_*j*_ is the corresponding integral weight, which is calculated based on Legendre polynomials and automatically computed by the Julia command `gausslegendre`. In Algorithm 2 we summarise the numerical recipe.

##### Algorithm 2

Generating-function-based parameter inference accounting for extrinsic noise and multiple gene copies

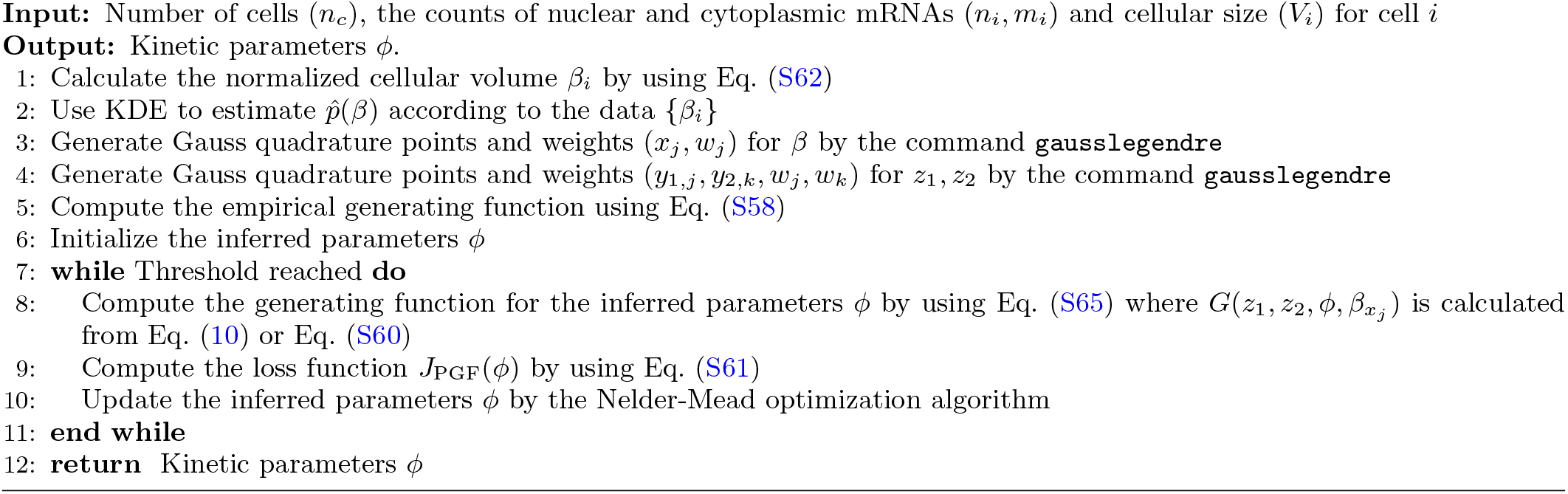

#### B. Finite-state-projection-based parameter inference

Given the *n*_*c*_ tuples of counts 𝒟= (*n*_*i*_, *m*_*i*_), the total likelihood of all observations is the product over every tuple, i.e.,

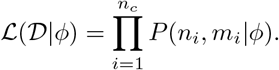

The inference of the kinetic parameters proceeds by minimizing the negative log likelihood, that is

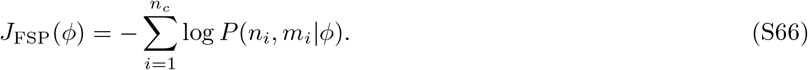

The optimization details of minimizing Eq. (S66) are the same as those of the inference method using the generating function. For Model I (Eq. (9)), we use finite state projection (FSP) to solve the joint probability for a single gene copy and for a given 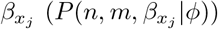 from Eqs. (S22), (S25) and (S27) in the steady state limit. The stationary distribution for the cell population is computed using

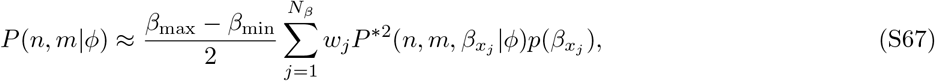

for the same tuple 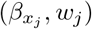 in Eq. (S65). The probability 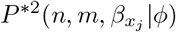 is the convolution of the probability 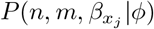. By plugging *P* (*n, m*|*ϕ*) into Eq. (S66), the negative log likelihood can be computed.

The numerical recipe for inference using FSP is summarised in Algorithm 3, and the optimization details are the same as those of the inference method using the generating function.

##### Algorithm 3

FSP-based parameter inference for Model I that accounts for extrinsic noise and multiple gene copies

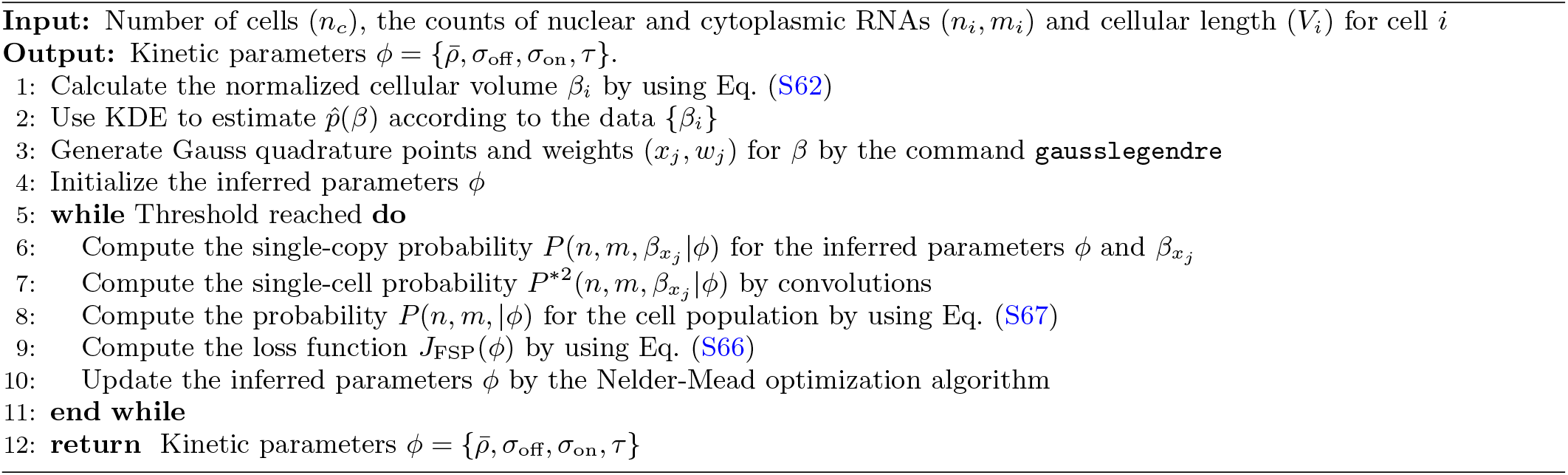

#### C. Method-of-moments-based parameter inference

We begin by deriving the first- and second-order moments for nuclear and cytoplasmic mRNA counts transcribed from two identical and independent gene copies in the presence of extrinsic noise. We here assume expression from each gene is described by Model I (9).

Let *n*_*i*_ and *m*_*i*_ denote the nuclear and cytoplasmic mRNA counts of gene copy *i* respectively. According to Eq. (S56), the corresponding means are

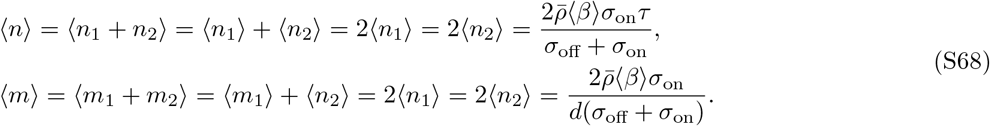

Applying the law of total variance var(*Y*) = ⟨var(*Y* |*Z*)⟩ + var(⟨*Y* |*Z*⟩), and the law of total covariance cov(*X, Y*) = ⟨cov(*X, Y* |*Z*)⟩ + cov(⟨*X*|*Z*⟩, ⟨*Y* |*Z*⟩), we find that the second-order moments are given by

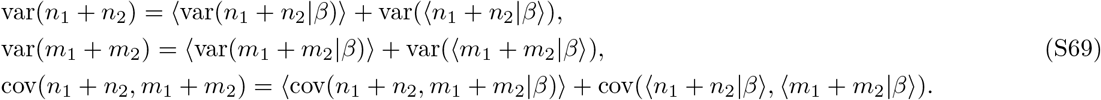

Since *n*_1_ and *n*_2_ are independent given a certain *β*, it follows that

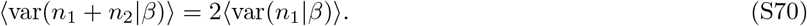

By similar arguments, we obtain that

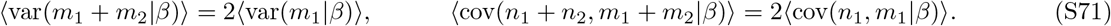

Next, using the linearity of the expectation operator, we find that

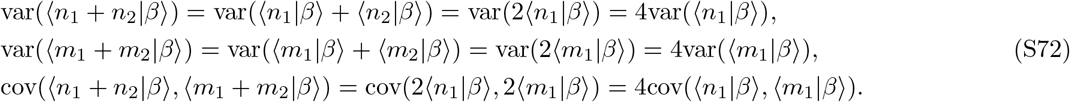

Combining Eqs. (S69)-(S72), we obtain

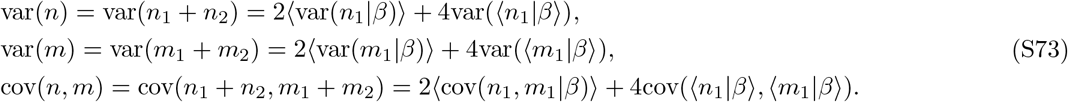

The terms on the RHS of Eq. (S73) are modified from those in Eqs. (S56) and (S57). These are given by

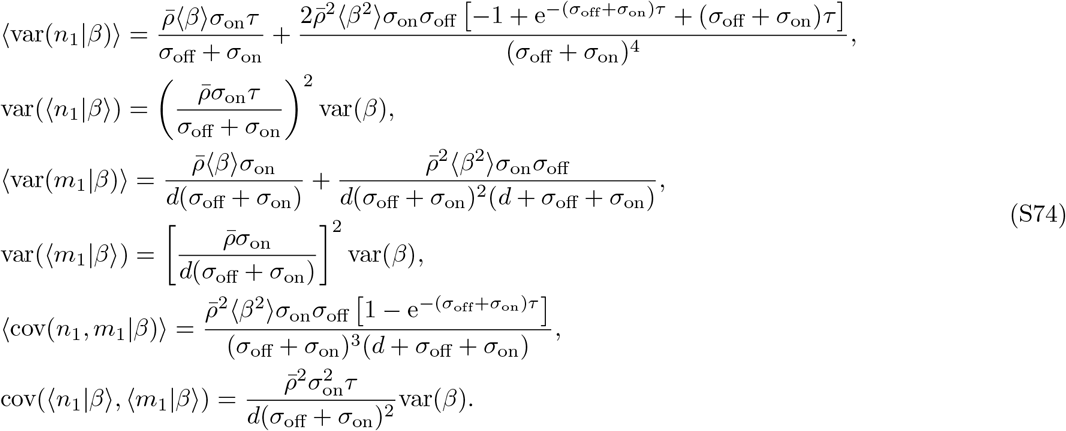

Parameter inference is performed by optimizing the loss function, which is given by

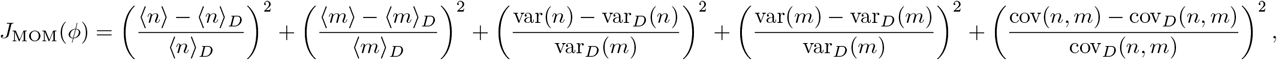

where the model’s moments are given by Eq. (S68), Eq. (S73) and Eq. (S74), while the moments with subscript *D* are those directly computed from data. This optimization problem is also solved by the Nelder-Mead algorithm.

### IX. Processing of MERFISH data

We sourced total mRNA counts and nuclear mRNA counts from the files `pnas.1912459116.sd12.csv` and `pnas.1912459116.sd14.csv` of Ref. [14]. There are 10,050 genes across a population of *n*_*c*_ = 1368 human osteosarcoma (U-2 OS) cells reported in the dataset. By excluding genes for which the total mRNA counts *<* 1, only 6724 genes remained. Afterwards, we processed the data from each gene into the tuples of nuclear-cytoplasmic mRNA counts (*n*_*i*_, *m*_*i*_) where *i* is the cell index. The number of tuples varied from gene to gene because for each gene we removed those cells whose nuclear or cytoplasmic counts were outliers; specifically from the data for gene *j*, we removed those cells whose counts for gene *j* were outside of the range (mean − 5 std, mean + 5 std) where the mean and standard deviation of counts for gene *j* are computed over the whole cell population.

### X. Generating-function-based model selection using 10-fold cross validation and ontology analysis

The purpose of model selection in this context is to identify the type of gene activity. Since the generating-function-based inference is not based on the conventional maximum likelihood method, it is difficult to see how any generating-function-based model selection method can be based on information criterion (such as the Akaike or Bayesian Information Criteria).

To avoid this difficulty, we propose to use 10-fold cross validation. We start by arranging candidate models in order of increasing complexity (number of parameters) and dividing the cell data into 10 equal-sized subsamples. Inference is repeated 10 times, where for each time, nine of the subsamples are used to infer parameters for candidate models, while the tenth is used to compute a performance score 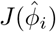 using Eq. (14). This process generates a score vector 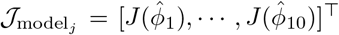 for each model *j* (this index increases with model complexity as described earlier). We denote the index of the proposed best model, i.e. that with the smallest mean score, by *j* = ℐ. The mean and standard deviation of this model’s score vector 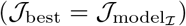 are denoted 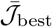 and *σ*_best_, respectively. To balance model complexity with fitting accuracy, we need to compare the proposed best model with simpler models having a smaller number of parameters. Defining 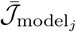 as the mean of the score vector for model *j*, for models with *j <*, ℐwe apply the inequality

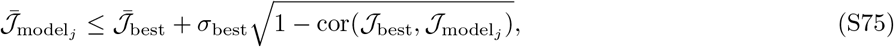

selecting the model with the smallest *j* that satisfies Eq. (S75) as the final best model. Note that cor(*x, y*) stands for the correlation coefficient of two score vectors *x* and *y*.

We summarise the entire model selection procedure in the following algorithm.

#### Algorithm 4

Generating-function-based model selection method using 10-fold cross validation

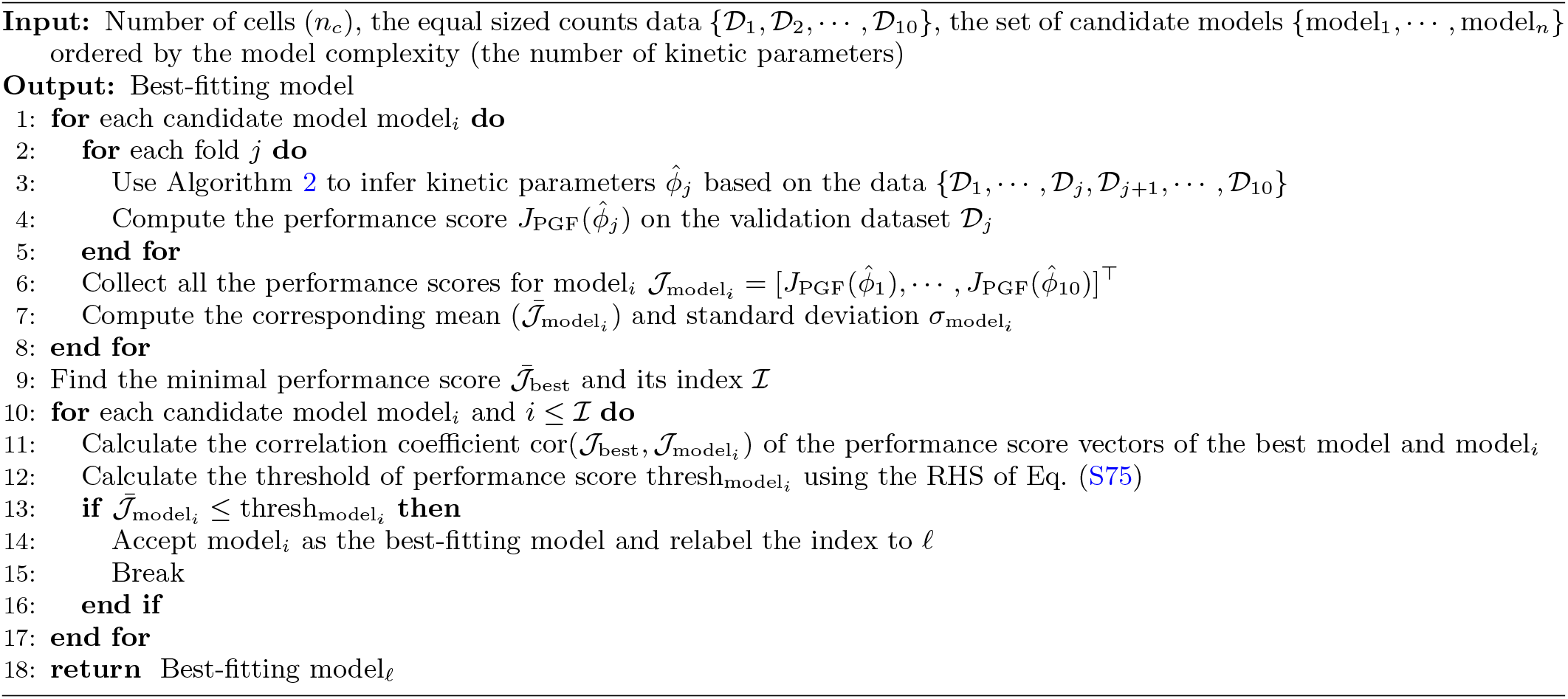

For the MERFISH data, we apply two candidate models – Poisson (Eq. (16)) and Model I (Eq. (9)) – to identify gene activity. The PGFs for a single gene copy without extrinsic noise are provided in Eqs. (S60), and (10), respectively. Using Eq. (S65) and these PGFs, we compute the PGF accounting for extrinsic noise and multiple gene copies, enabling the application of Algorithm 4. Notably, we exclude the refractory model (Model II) as a candidate model due to its low accuracy in being correctly selected when using sample sizes of ∼ 10^3^ 10^4^ cells (see Fig. S4). Given the genes classified according to one of 2 best fitting models, subsequently we utilized the DAVID software [57] for gene ontology (GO) analysis of each of the 2 gene categories. The analysis was contextualized against a background of filtered 6724 genes. GO terms were extracted from the `GOTERM_BP_FAT` category. Our criteria for inclusion were a Benjamini *p*-value (*p*_adj_) of less than 0.01 and a minimum of 15 enriched genes per term. These terms were then ranked based on Gene Ratio. The comprehensive results are detailed in Excel files stored in Github.

### XI. Code and data availability

All codes and data have been deposited at https://github.com/edwardcao3026/pgf-inf.

**FIG. S1.**
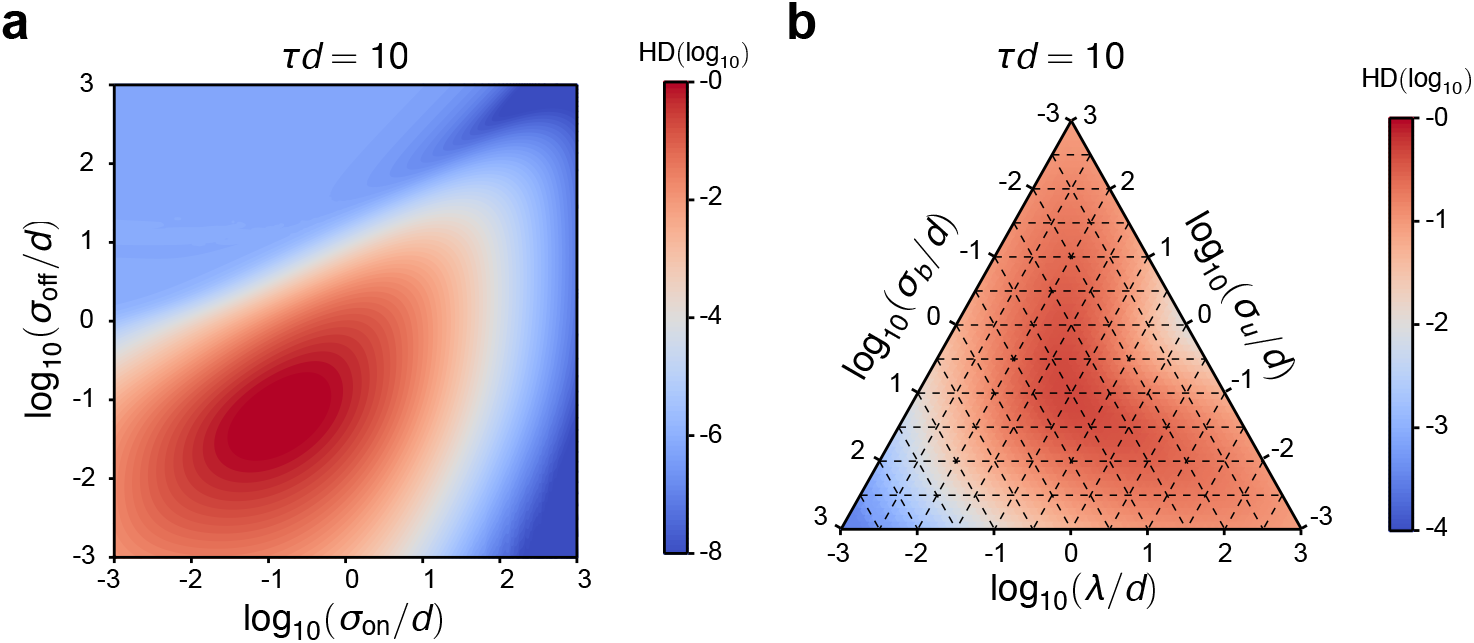
Hellinger distance (HD) between the joint distributions of non-Markovian models (Models I and II) and Markovian Models (Model III and IV) as a function of the normalised gene state switching rates *σ*_on_/*d* and *σ*_off_/*d* (or *σ*_u_/*d, σ*_b_/*d* and *λ/d*), and the normalised export time *τd* = 10 (*ρ* is fixed to 5). (a) Comparison of Model I vs Model III; (b) Comparison of Model II vs Model IV.

**FIG. S2.**
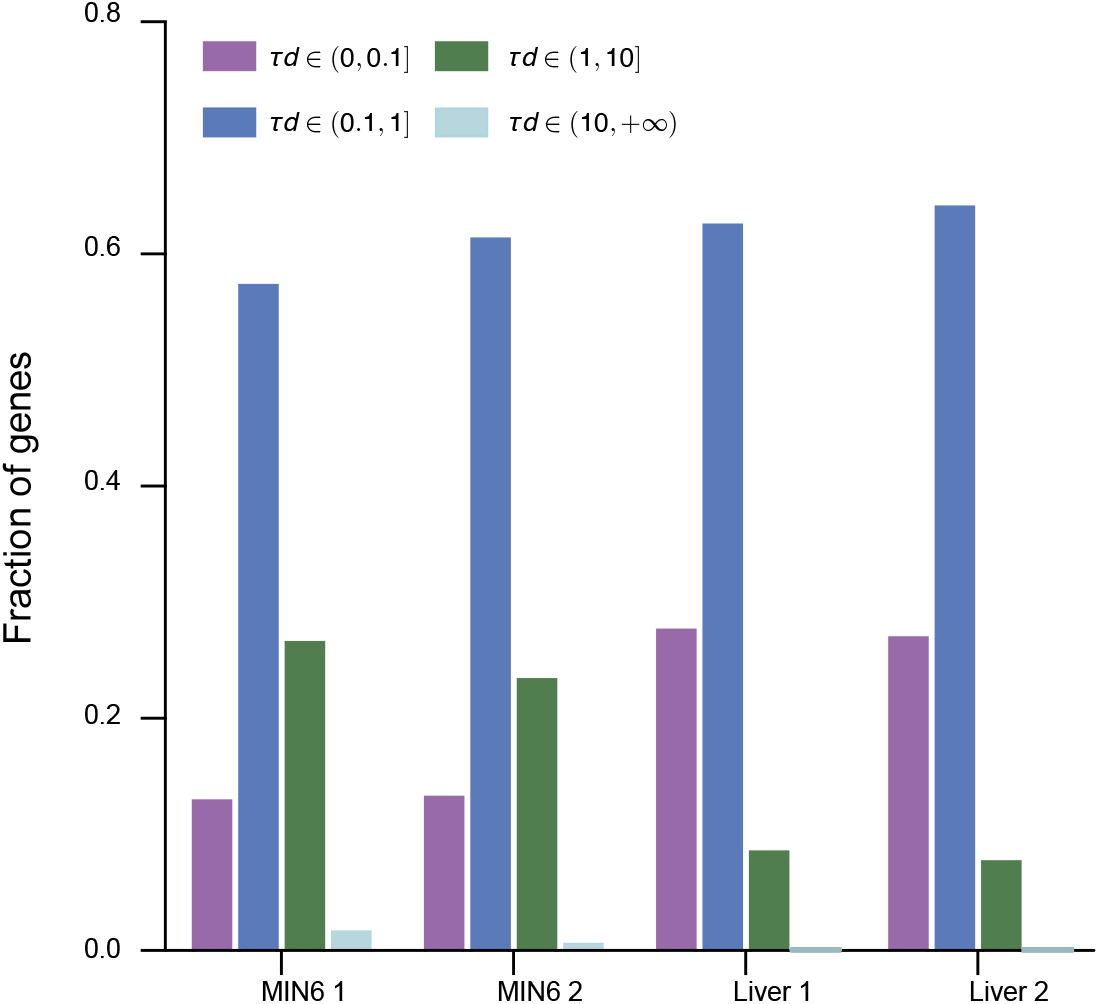
Distributions of *τd* estimated from mRNA data measured in two samples of MIN6 cells and liver cells.

**FIG. S3.**
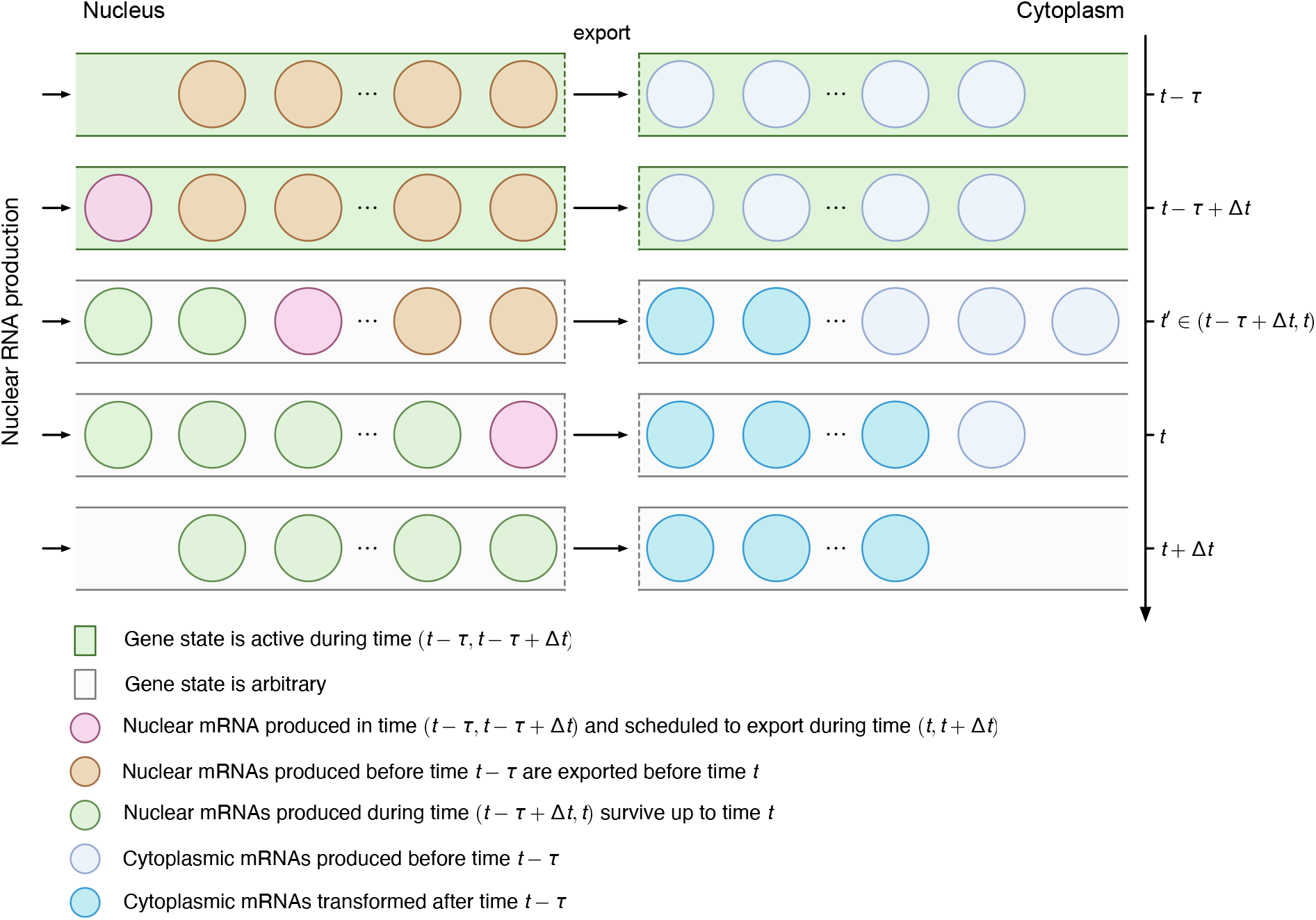
Illustration of the chain of events culminating in the export of a nuclear mRNA in Model I (refer to event (C)). Let’s assume there are *n*′ nuclear mRNAs (brown balls) and *m*′ cytoplasmic mRNAs (purple balls) produced before time *t* − *τ*. In the next time step *t* −*τ* + Δ*t*, there is a newborn nuclear mRNA (red ball). During (*t* − *τ* + Δ*t, t*), the first *n*′ brown mRNAs are progressively exported and turned into cytoplasmic mRNAs (blue balls) and in the meanwhile *n* new nuclear mRNAs (green balls) are produced. Since each nuclear RNA stays for a fixed time *τ*, the red RNA definitely leaves in the time interval (*t, t* + Δ*t*). Note that during the time interval (*t* − *τ* + Δ*t, t*), the red RNA does not participate in any dynamics of the system which explains Eq. (S15) and the dynamics of the green mRNAs follows Eq. (S22) which is a simple telegraph model without decay from (0, *τ*).

**FIG. S4.**
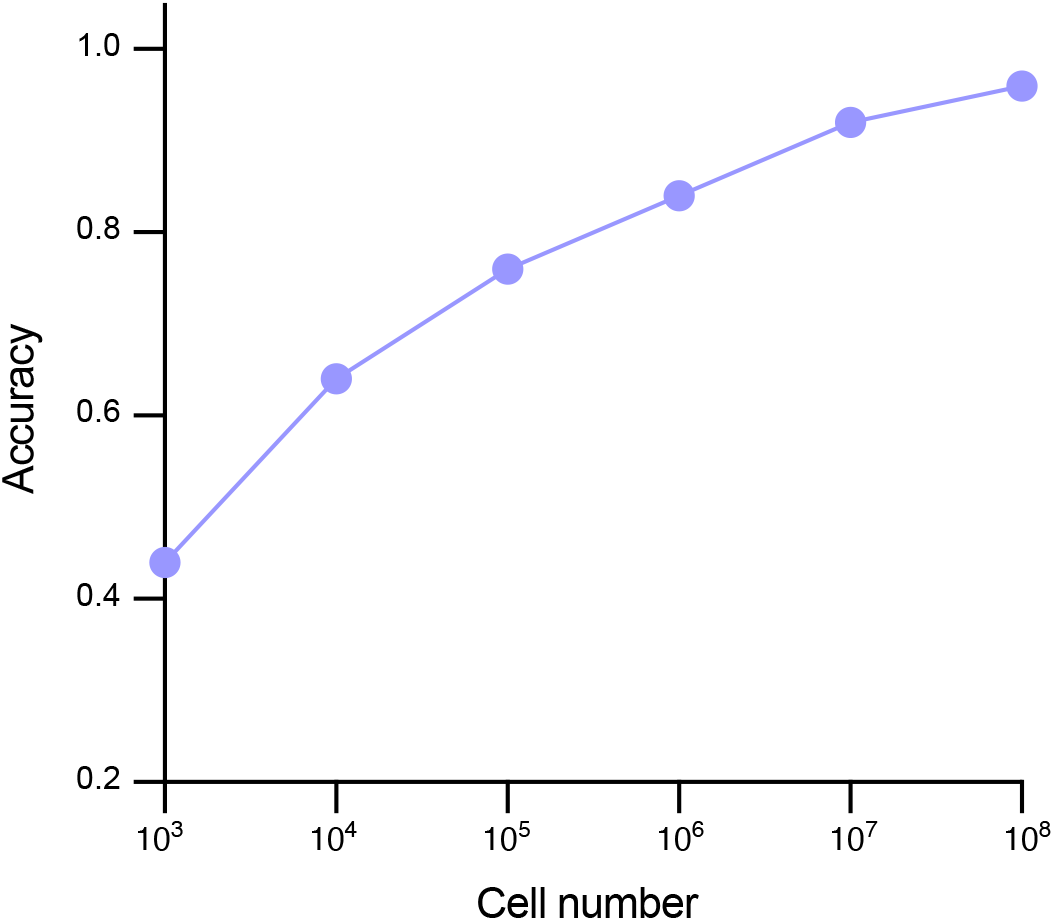
Accuracy of identifying the true model (Model II, Eq. (12)) as a function of the number of cells in the sample (*n*_*c*_). The counts data are generated by simulating Model II using the delay SSA. The kinetic parameters are shown in Table S4. The model selection procedure in SM X is applied to the data to select between two candidate models: Models I and II. The accuracy of identifying the correct model increases with the number of cells. However, in a typical sample size *n*_*c*_ = 10^3^ −10^4^, the two competing models are difficult to discern. Note that for simplicity, here we assumed that all noise is intrinsic and there is a single gene copy; a similar outcome is expected in the presence of extrinsic noise and for multiple gene copies.

**FIG. S5.**
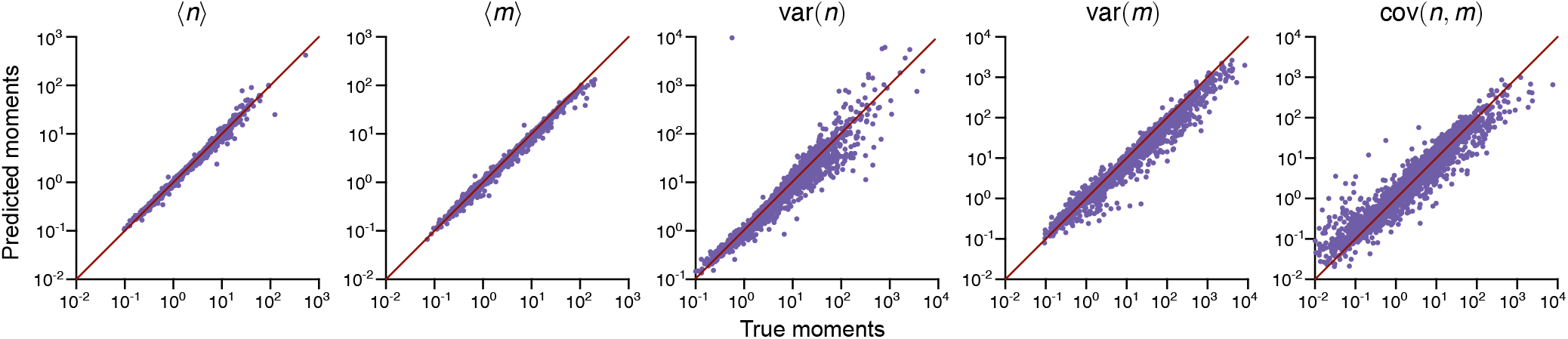
Accuracy of the first- and second-order moments predicted by fitting Model I to the mRNA count data from 2,722 non-Poissonian genes in the MERFISH dataset. The moments are calculated using Eqs. (S68), (S73), and (S74) and compared with the true moments from experimental data. 2,184 genes meet the criterion that the average relative error across the five moments is less than 35%.

**TABLE S1.**
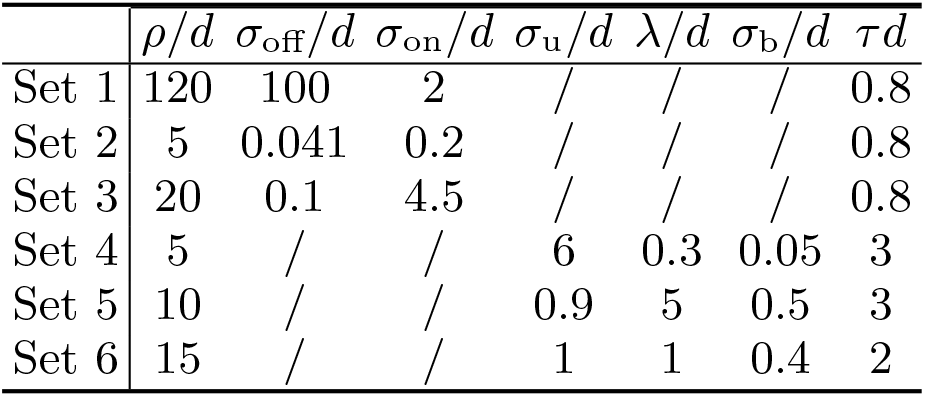
Kinetic parameters used in the P-P plots of Fig. 2a. The degradation rate *d* is set to 1.

| | $\rho/d$ | $\sigma_{\text{off}}/d$ | $\sigma_{\text{on}}/d$ | $\sigma_{\text{u}}/d$ | $\lambda/d$ | $\sigma_{\text{b}}/d$ | $\tau d$ |
| --- | --- | --- | --- | --- | --- | --- | --- |
| Set 1 | 120 | 100 | 2 | / | / | / | 0.8 |
| Set 2 | 5 | 0.041 | 0.2 | / | / | / | 0.8 |
| Set 3 | 20 | 0.1 | 4.5 | / | / | / | 0.8 |
| Set 4 | 5 | / | / | 6 | 0.3 | 0.05 | 3 |
| Set 5 | 10 | / | / | 0.9 | 5 | 0.5 | 3 |
| Set 6 | 15 | / | / | 1 | 1 | 0.4 | 2 |

**TABLE S2.** Inferred and true kinetic parameters from simulated data – data summarised in Fig. 3b. The true parameters are uniformly sampled from 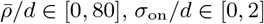 and *σ*_off_/*d* ∈ [0, 40], – these are roughly in the range estimated for various types of eukaryotic cells in Table I of Ref. [58]. The parameter *dτ* is sampled from *dτ* ∈ [0, 1] since about 80% of genes fall in this range according to Fig. S2.

|  | PGF |  |  |  | FSP |  |  |  | True |  |  |  |
| --- | --- | --- | --- | --- | --- | --- | --- | --- | --- | --- | --- | --- |
| | $\bar{\rho}/d$ | $\sigma_{\text{off}}/d$ | $\sigma_{\text{on}}/d$ | $\tau d$ | $\bar{\rho}/d$ | $\sigma_{\text{off}}/d$ | $\sigma_{\text{on}}/d$ | $\tau d$ | $\bar{\rho}/d$ | $\sigma_{\text{off}}/d$ | $\sigma_{\text{on}}/d$ | $\tau d$ |
| Set 1 | 57.46 | 24.31 | 2.01 | 0.14 | 69.08 | 28.07 | 1.91 | 0.15 | 60.97 | 24.71 | 1.94 | 0.15 |
| Set 2 | 29.19 | 20.53 | 0.49 | 0.66 | 24.75 | 19.33 | 0.53 | 0.63 | 27.14 | 19.48 | 0.52 | 0.61 |
| Set 3 | 36.53 | 9.51 | 0.1 | 0.28 | 36.64 | 7.05 | 0.08 | 0.34 | 40.62 | 8.84 | 0.09 | 0.32 |
| Set 4 | 74.48 | 37.82 | 0.92 | 0.26 | 52.06 | 22.88 | 0.85 | 0.27 | 73.26 | 30.41 | 0.79 | 0.28 |
| Set 5 | 24.29 | 13.17 | 0.25 | 0.57 | 18.36 | 8.69 | 0.23 | 0.58 | 25.01 | 11.87 | 0.22 | 0.57 |
| Set 6 | 3.90 | 0.11 | 0.38 | 0.60 | 3.79 | 0.08 | 0.32 | 0.59 | 4.06 | 0.16 | 0.42 | 0.61 |
| Set 7 | 39.73 | 22.60 | 1.41 | 0.20 | 58.11 | 40.48 | 1.61 | 0.18 | 44.07 | 28.13 | 1.43 | 0.18 |
| Set 8 | 19.28 | 32.81 | 1.2 | 0.8 | 9.35 | 14.02 | 1.12 | 0.8 | 18.53 | 31.42 | 1.18 | 0.82 |
| Set 9 | 51.12 | 19.68 | 1.42 | 0.5 | 49.9 | 18.51 | 1.38 | 0.51 | 51.13 | 19.59 | 1.42 | 0.51 |
| Set 10 | 35.36 | 4.56 | 1.12 | 0.26 | 41.34 | 5.41 | 1.45 | 0.28 | 35.34 | 4.59 | 1.08 | 0.26 |
| Set 11 | 23.23 | 9.37 | 0.56 | 0.64 | 23.94 | 11.58 | 0.62 | 0.62 | 26.67 | 12.13 | 0.61 | 0.58 |
| Set 12 | 4.52 | 0.25 | 0.19 | 0.58 | 4.26 | 0.22 | 0.18 | 0.57 | 4.44 | 0.23 | 0.17 | 0.57 |
| Set 13 | 32.28 | 16.63 | 0.40 | 0.73 | 25.91 | 14.09 | 0.42 | 0.67 | 27.14 | 15.04 | 0.43 | 0.61 |
| Set 14 | 26.11 | 7.50 | 0.35 | 0.19 | 32.86 | 9.29 | 0.35 | 0.20 | 39.14 | 10.04 | 0.31 | 0.17 |
| Set 15 | 49.8 | 21.16 | 1.55 | 0.38 | 46.11 | 19.66 | 1.56 | 0.38 | 47.25 | 20.94 | 1.61 | 0.38 |

**TABLE S3.** Inferred and true kinetic parameters and extrinsic nose from simulated data – data summarised in Fig. 3c. The true parameters are uniformly sampled from 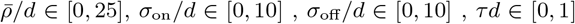. The extrinsic noise is sampled as *β* ∈Gamma(40, 0.025).

|  | Joint |  |  |  | Moments |  |  |  | True |  |  |  |
| --- | --- | --- | --- | --- | --- | --- | --- | --- | --- | --- | --- | --- |
| | $\bar{\rho}/d$ | $\sigma_{\text{off}}/d$ | $\sigma_{\text{on}}/d$ | $\tau d$ | $\bar{\rho}/d$ | $\sigma_{\text{off}}/d$ | $\sigma_{\text{on}}/d$ | $\tau d$ | $\bar{\rho}/d$ | $\sigma_{\text{off}}/d$ | $\sigma_{\text{on}}/d$ | $\tau d$ |
| Set 1 | 7.132 | 3.838 | 2.041 | 0.745 | 6.644 | 2.922 | 1.743 | 0.746 | 7.1 | 3.67 | 1.98 | 0.711 |
| Set 2 | 18.211 | 1.195 | 6.462 | 0.28 | 22.445 | 3.96 | 16.384 | 0.277 | 17.915 | 1.677 | 9.08 | 0.282 |
| Set 3 | 14.508 | 1.256 | 4.792 | 0.342 | 13.46 | 0.539 | 2.859 | 0.359 | 14.225 | 1.06 | 4.739 | 0.344 |
| Set 4 | 14.095 | 6.335 | 3.533 | 0.461 | 12.864 | 4.631 | 2.919 | 0.478 | 15.531 | 6.961 | 3.29 | 0.48 |
| Set 5 | 25.567 | 2.773 | 2.825 | 0.986 | 26.083 | 3.441 | 3.303 | 0.987 | 24.335 | 2.45 | 2.812 | 0.956 |
| Set 6 | 23.553 | 5.498 | 1.056 | 0.506 | 21.522 | 5.004 | 1.057 | 0.503 | 22.538 | 5.162 | 1.046 | 0.494 |
| Set 7 | 18.991 | 1.334 | 6.041 | 0.276 | 17.535 | 0.486 | 3.243 | 0.29 | 19.29 | 1.651 | 6.965 | 0.275 |
| Set 8 | 4.792 | 10.121 | 2.1 | 0.495 | 4.949 | 9.467 | 1.874 | 0.514 | 4.472 | 9.755 | 2.244 | 0.502 |
| Set 9 | 11.499 | 4.763 | 2.836 | 0.553 | 10.97 | 4.835 | 3.067 | 0.552 | 11.128 | 4.812 | 2.983 | 0.558 |
| Set 10 | 4.207 | 7.81 | 4.273 | 0.202 | 2.988 | 3.328 | 3.199 | 0.201 | 3.895 | 8.666 | 5.451 | 0.186 |
| Set 11 | 8.45 | 1.37 | 0.9 | 0.679 | 8.498 | 1.394 | 0.885 | 0.685 | 8.469 | 1.4 | 0.898 | 0.656 |
| Set 12 | 8.174 | 4.189 | 5.198 | 0.383 | 7.597 | 2.713 | 3.785 | 0.416 | 7.904 | 3.399 | 4.411 | 0.391 |
| Set 13 | 7.88 | 4.614 | 1.101 | 0.529 | 8.743 | 5.835 | 1.254 | 0.504 | 7.555 | 4.562 | 1.128 | 0.527 |
| Set 14 | 6.257 | 0.187 | 1.606 | 0.421 | 6.423 | 0.377 | 2.402 | 0.429 | 6.494 | 0.389 | 2.457 | 0.417 |
| Set 15 | 1.273 | 1.624 | 0.737 | 0.559 | 1.344 | 1.618 | 0.733 | 0.517 | 1.531 | 2.253 | 0.723 | 0.559 |

**TABLE S4.** The sets of kinetic parameters used for generating counts data for model selection in Fig. S4. These parameters are sampled from *ρ/d* ∈ [0, 30], *σ*_u_/*d* ∈ [0, 10], *λ/d* ∈ [0, 10], *σ*_b_/*d* ∈ [0, 10], *τd* ∈ [0, 5] and *d* = 1.

|  | Parameters |  |  |  |  |
| --- | --- | --- | --- | --- | --- |
| | $\rho/d$ | $\sigma_u/d$ | $\lambda/d$ | $\sigma_b/d$ | $\tau d$ |
| Set 1 | 7.218 | 5.533 | 7.089 | 7.148 | 0.286 |
| Set 2 | 6.575 | 2.567 | 4.518 | 8.341 | 4.598 |
| Set 3 | 8.329 | 4.982 | 4.374 | 5.063 | 4.564 |
| Set 4 | 9.487 | 6.055 | 6.067 | 6.366 | 3.178 |
| Set 5 | 13.404 | 5.804 | 1.592 | 8.433 | 3.625 |
| Set 6 | 15.858 | 1.573 | 7.57 | 9.463 | 1.914 |
| Set 7 | 10.974 | 4.815 | 5.871 | 3.92 | 2.05 |
| Set 8 | 27.07 | 3.895 | 4.48 | 6.472 | 2.668 |
| Set 9 | 2.453 | 3.566 | 8.616 | 5.964 | 3.893 |
| Set 10 | 14.149 | 5.464 | 1.975 | 9.687 | 4.062 |
| Set 11 | 0.675 | 9.511 | 2.835 | 5.225 | 3.558 |
| Set 12 | 28.847 | 7.508 | 6.949 | 2.481 | 4.648 |
| Set 13 | 3.022 | 1.967 | 9.764 | 1.52 | 0.777 |
| Set 14 | 1.463 | 0.597 | 9.872 | 1.859 | 0.245 |
| Set 15 | 9.632 | 8.053 | 7.964 | 2.949 | 0.092 |
| Set 16 | 12.865 | 5.673 | 5.804 | 4.397 | 2.641 |
| Set 17 | 8.296 | 2.601 | 0.968 | 7.805 | 4.486 |
| Set 18 | 19.2 | 9.526 | 5.221 | 0.445 | 2.083 |
| Set 19 | 5.62 | 5.143 | 1.069 | 9.366 | 1.774 |
| Set 20 | 2.545 | 9.837 | 3.254 | 3.468 | 2.365 |
| Set 21 | 14.819 | 1.984 | 0.524 | 7.203 | 0.11 |
| Set 22 | 16.171 | 3.753 | 2.294 | 2.984 | 0.889 |
| Set 23 | 16.479 | 2.499 | 9.822 | 4.521 | 0.796 |
| Set 24 | 24.963 | 8.934 | 7.695 | 0.67 | 0.69 |
| Set 25 | 19.399 | 8.837 | 9.969 | 5.244 | 0.724 |

**TABLE S5.** Dependence of the results of the model selection and parameter inference procedure on various assumptions and the type of input single-cell data. The first column shows whether the model accounts for both intrinsic and extrinsic noise (tick) or only intrinsic noise (cross). The second column is the assumed number of gene copies; for eukaryotic genes, this can be 2 or 4 depending on the position in the cell-cycle (information that is not available for the MERFISH dataset). The third column shows the type of data used as input to the inference algorithm: total counts (sum of nuclear and cytoplasmic counts) or joint data. The fourth, fifth and sixth columns show the medians of the estimated *f*_on_, burst size and burst frequency for Model I. The number of genes that are best fit by the Poisson model and Model I are shown in columns 7 and 8, respectively. The number of non-Poissonian genes that pass the quality criteria (similar to Fig. S5) are shown in the last column; the parameters for Model I are only estimated for these genes. When performing inference on total count data, we set *z*_1_ = 1 in Eq. (10); this reduces the generating function solution of Model I to that of the conventional telegraph model [12]. This choice conforms with the common practice of interpreting the mRNA count prediction of the telegraph model as the total mRNA per cell. For experimental sets where extrinsic noise was not considered, we fixed *β* = 1 in Algorithm 2. In experimental sets involving four gene copies, we set the exponent of *G* in Eq. (S65) to 4.

| | Extrinsic noise | Gene copy numbers | Data type | $f_{\text{on}}$ | Burst size | Burst frequency | # of Poissonian genes | # of non-Poissonian genes | # of non-Poissonian genes passing fitting quality check |
| --- | --- | --- | --- | --- | --- | --- | --- | --- | --- |
| Set 1 | × | 2 | Joint | 0.66 | 7.84 | 0.59 | 617 | 6107 | 4811 |
| Set 2 | × | 2 | Total | 0.19 | 3.05 | 1.59 | 626 | 6098 | 6056 |
| Set 3 | ✓ | 2 | Joint | 0.64 | 6.83 | 0.69 | 4022 | 2722 | 2184 |
| Set 4 | ✓ | 2 | Total | 0.40 | 5.41 | 1.14 | 4172 | 2552 | 2523 |
| Set 5 | × | 4 | Joint | 0.50 | 6.52 | 0.29 | 624 | 6100 | 5389 |
| Set 6 | × | 4 | Total | 0.15 | 3.10 | 0.76 | 622 | 6102 | 6069 |
| Set 7 | ✓ | 4 | Joint | 0.44 | 3.88 | 0.39 | 3987 | 2737 | 2241 |
| Set 8 | ✓ | 4 | Total | 0.34 | 6.35 | 0.48 | 4169 | 2555 | 2527 |

